# Mechanism of *in vivo* activation of the MutLγ-Exo1 complex for meiotic crossover formation

**DOI:** 10.1101/2019.12.16.876623

**Authors:** Aurore Sanchez, Céline Adam, Felix Rauh, Yann Duroc, Lepakshi Ranjha, Bérangère Lombard, Xiaojing Mu, Damarys Loew, Scott Keeney, Petr Cejka, Raphaël Guérois, Franz Klein, Jean-Baptiste Charbonnier, Valérie Borde

## Abstract

Crossovers generated during the repair of programmed double-strand breaks (DSBs) during homologous recombination are essential for fertility to allow accurate homolog segregation during the first meiotic division. Most crossovers arise through the cleavage of recombination intermediates by the Mlh1-Mlh3 (MutLγ) endonuclease and an elusive non-catalytic function of Exo1, and require the Polo kinase Cdc5. Here we show in budding yeast that MutLγ forms a constitutive complex with Exo1, and in meiotic cells transiently contacts the Msh4-Msh5 (MutSγ) heterodimer, also required for crossover formation. We further show that MutLγ-Exo1 associates with recombination intermediates once they are committed to the crossover repair pathway, and then Exo1 recruits Cdc5 through a direct interaction that is required for activating MutLγ and crossover formation. Exo1 therefore serves as a non-catalytic matchmaker between Cdc5 and MutLγ. We finally show that *in vivo*, MutLγ associates with the vast majority of DSB hotspots, but at a lower frequency near centromeres, consistent with a strategy to reduce at-risk crossover events in these regions. Our data highlight the tight temporal and spatial control of the activity of a constitutive, potentially harmful, nuclease.

## Introduction

Meiotic recombination provides crossovers that allow homologous chromosomes to be transiently physically attached together and then properly segregated^1^. A lack or altered distribution of meiotic crossovers is a major source of aneuploidies, leading to sterility or disorders such as the Down syndrome, highlighting the importance of understanding how crossovers are controlled during meiosis. Meiotic recombination is triggered by the formation of programmed DNA DSBs, catalyzed by Spo11 together with several conserved protein partners^2,3^. Following Spo11 removal, 5’ DSB ends are resected and the 3’ ends invade a homologous template, leading to formation of D-loop intermediates. After DNA synthesis, many of these intermediates are dismantled by helicases and repaired without a crossover^4^. However, a subset are stabilized, and upon capture of the second DSB end, mature into a double Holliday junction (dHJ), which is almost exclusively resolved asymmetrically as a crossover^5,6^. Essential to this crossover pathway are a group of proteins, called ZMMs, which collectively stabilize and protect recombination intermediates against helicases^4,7–10^. The mismatch repair MutLγ (Mlh1-Mlh3) heterodimer is proposed to subsequently act on these ZMM-stabilized dHJs to resolve them into crossovers^11^. Accordingly, MutLγ forms foci on mammalian and plant meiotic chromosomes in numbers and distribution that mirror those of crossovers (reviewed in^12^).

In eukaryotic mismatch repair, a MutS-related complex (MutSα or MutSβ) recognizes a mismatch, and recruits a MutL-related complex, to initiate repair together with PCNA and Exo1 (reviewed in^13^). MutLα represents the major mismatch repair activity, which involves its endonuclease activity^14–16^. By contrast, MutLγ is less important for mismatch repair, but has an endonuclease activity essential for its function in meiotic crossovers^11,17–23^. The mechanism of dHJ resolution and crossover formation by MutLγ is still unknown. MutLγ alone does not resolve HJs *in vitro*^20,21,24^, and may need additional partners, meiosis-specific post-translational modifications and/or a specific DNA substrate to promote specific nuclease activity and hence crossover formation.

Based on recent *in vitro* experiments and genome-wide analysis of meiotic recombination, it seems that rather than acting as a canonical resolvase, MutLγ may nick the DNA, which could result in dHJ resolution and crossover formation if two closely spaced nicks on opposite strands are made^24,25^. *In vitro* analyses with recombinant proteins also suggested that MutLγ may need to extensively polymerize along several kilobases of DNA in order to activate its DNA cleavage activity^24^.

MutLγ partners with MutSγ (Msh4-Msh5), part of the ZMM complex proposed to first stabilize recombination intermediates and then to work together with MutLγ for crossover resolution (reviewed in^10,12,13^). In addition, the exonuclease Exo1 interacts *in vitro* with Mlh1. This interaction with Mlh1, but not Exo1 catalytic activity, is essential for MutLγ function in crossover formation^26,27^. This is in contrast with MMR, where Exo1’s catalytic activity is not absolutely required because of redundant exonuclease activities *in vivo*^28^. Exo1’s meiotic crossover function appears to be conserved because *Exo1 null* mice are sterile and show reduced crossovers, whereas catalytic mutants of Exo1 are fertile^29,30^. How Exo1 is required for activating MutLγ is not known. Finally, the polo-like kinase Cdc5 kinase is required for MutLγ activity in meiotic crossover^31^. Cdc5 expression is induced by the Ndt80 transcription factor at the pachytene stage of meiotic prophase, once homolog synapsis and recombination intermediates formation are achieved^32–35^. Cdc5 promotes multiple steps during exit from pachytene, including dHJ resolution, disassembly of the synaptonemal complex, and sister kinetochore mono-orientation^36–38^. Among other targets, Cdc5 activates the Mus81-Mms4 structure-specific nuclease, a minor crossover activity in meiosis, by phosphorylating Mms4^39^. However, it is not known how Cdc5 promotes MutLγ’s major crossover activity.

We undertook a thorough analysis of MutLγ activation and distribution genome-wide in budding yeast. We found that MutLγ and Exo1 form a constitutive complex that associates on recombination sites stabilized by ZMM proteins. Exo1 then serves as a recruiting platform for the Cdc5 kinase through a direct, specific interaction that is necessary for crossover formation by MutLγ. Finally, we find that MutLγ-dependent crossover formation is regulated by chromosome structure at multiple levels.

## Results

### MutLγ forms a complex with Exo1 and transiently interacts with MutSγ *in vivo*

A recent crystal structure of the C-terminal domain (Cter) of MutLγ shows that the last residues of Mlh1 are part of the Mlh3 endonuclease active site, rendering it impossible to C-terminally tag Mlh1 (Figure 1A) (Dai et al, in preparation)^40^. To preserve MutLγ functionality *in vivo*, we introduced an internal tag at an unstructured loop in Mlh3 that is predicted not to affect MutLγ function and catalytic activity (Figure 1A). Tags inserted at this site had no detectable effect on mismatch repair, spore viability or meiotic crossover frequency, and are therefore suitable for further *in vivo* studies (Figure S1).

**Figure 1:**
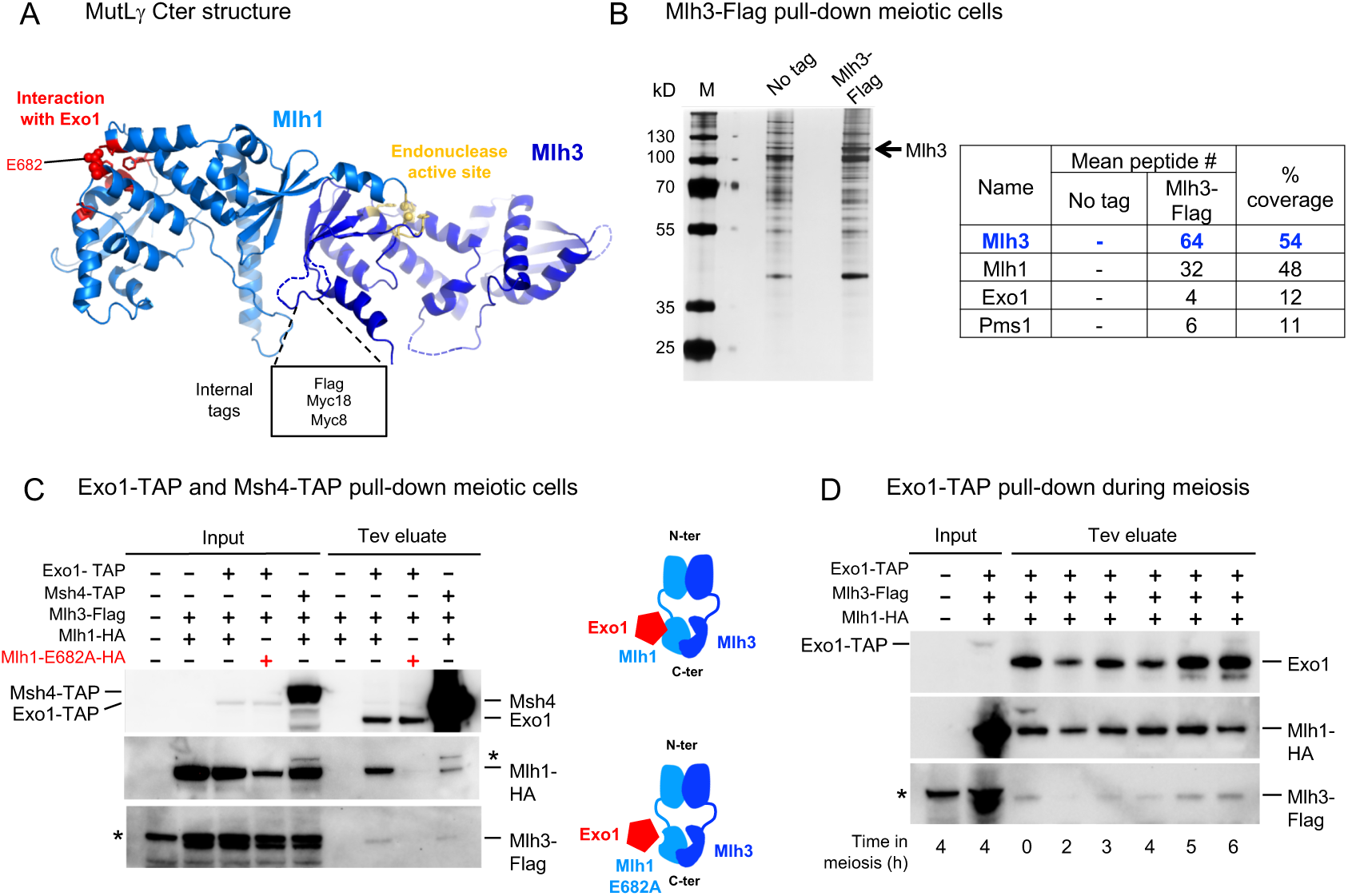
MutLγ forms a complex with Exo1 and transiently interacts with MutSγ *in vivo* (A) Crystal structure of the C-terminal region of *S. cerevisiae* Mlh1-Mlh3 heterodimer (Dai et al, in preparation). The Mlh1 and Mlh3 regions are colored in light and dark blue, respectively. The Mlh1 binding motif for Exo1 and the endonuclease site of Mlh3 are colored in red and yellow, respectively. (B) Affinity pull-down of Mlh3-Flag from cells at 4 h in meiosis. Left: silver-stained gel of pulled-down proteins. Right: mass-spectrometry analysis of selected proteins reproducibly identified in all replicates and not in the controls (no tag strain). The number of peptides of 4 independent experiments is shown (two with benzonase treatment, two without). Detail of pulled-down proteins in Table S1. (C) Coimmunoprecipitation by Exo1-TAP or Msh4-TAP from cells at 4 h in meiosis analyzed by Western blot. (D) Coimmunoprecipitation by Exo1-TAP at the indicated times in meiosis. Asterisks indicate non-specific cross-hybridizing bands. See also Figure S1.

To ask if MutLγ acts in complex with other partners, we immunoprecipitated Mlh3 from cells at 4 h in meiosis, when crossovers appear^6^, followed by mass spectrometry analysis (Figure 1B). In four independent experiments, Mlh3 pulled down its MutLγ partner Mlh1, as expected, but also Exo1, in agreement with its proposed function to assist MutLγ for crossover resolution^27^ (Figure 1B and Table S1). We confirmed this interaction by Western blot analysis of the reverse pull down of Mlh3 by Exo1 (Figure 1C). Interestingly, Exo1 pulled down Mlh1 and Mlh3 throughout meiosis, suggesting that Exo1 is a constitutive partner of MutLγ (Figure 1D). This interaction required the Mlh1 E682 residue that mediates the direct interaction between Exo1 and Mlh1, suggesting that the interaction between Exo1 and MutLγ is mediated by Mlh1, as it is for Exo1 and MutLα during MMR^26^ (Figure 1C). Consistent with these results, an *mlh1E682A* mutant displays meiotic crossover defects^27^.

Surprisingly, we did not recover any peptides from the Msh4-Msh5 heterodimer or from other ZMM proteins in the Mlh3 immunoprecipitates (Table S1). A weak coimmunoprecipitation of Mlh1 and Mlh3 by Msh4 was detected, but the fraction of Mlh1 and Mlh3 was very low (Figure 1C). Altogether, these results indicate that MutLγ forms a constitutive complex with Exo1, distinct from the other ZMM proteins, and that it likely establishes only transient contacts with MutSγ.

### Mlh3 foci on yeast meiotic chromosomes are distinct from ZMM foci

As in plants and mammals, Mlh3 formed foci on yeast meiotic chromosomes (Figure 2A-D). Importantly, Mlh3 foci numbers were almost null in *zip3Δ* and substantially reduced in *msh4Δ* mutants, in agreement with genetic data showing that MutLγ acts downstream of ZMM proteins (Figures 2E, F and H). Surprisingly, although Zip3 and to a lesser extent Msh4 are important for Mlh3 foci, Mlh3 showed very little colocalization with either Zip3 or Msh4, suggesting that Mlh3 coexists only briefly with the other two components at the same intermediates (Figures 2A, 2B, and 2I). We conclude that Zip3 and Msh4 ZMM are required for normal Mlh3 foci formation, and propose that Zip3 and Msh4 form earlier foci on recombination intermediates, before Mlh3 binding, which may reflect their function for intermediates stabilization. Then, taking into account the interaction detected between MutLγ and Msh4 (Figure 1C), we propose that a fraction of Msh4 persists to play a later function with MutLγ.

**Figure 2:**
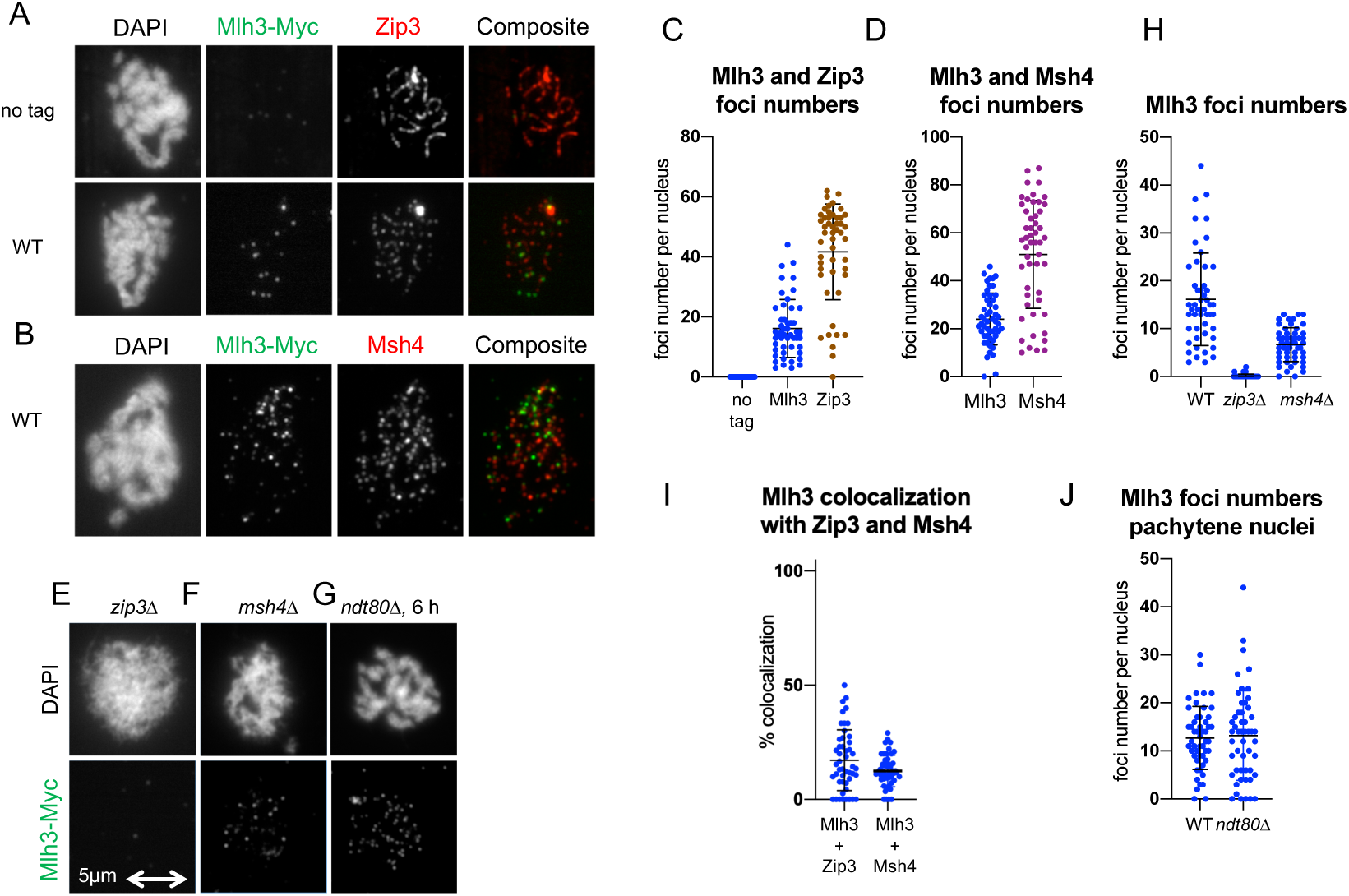
Mlh3 forms foci on yeast pachytene meiotic chromosomes, distinct from ZMM foci (A) Comparison of Mlh3-Myc18 and Zip3 foci. (B) Comparison of Mlh3-Myc18 and Msh4-HA foci. (C) and (D) Quantification of Mlh3, Zip3 and Msh4 foci from (A) and (B). (E) to (G): Mlh3-myc foci in *zip3Δ* (E), *msh4Δ* (F) and *ndt80Δ* (G). (H) Quantification of Mlh3-Myc foci in *zip3Δ* and *msh4Δ* mutants from (A), (E) and (F). 51 nuclei each examined. (I) Co-localization quantification of Mlh3 with Zip3 or Msh4 foci. The % of Mlh3 foci co-localizing is indicated (J) Quantification of Mlh3 foci in pachytene-selected nuclei (same strains as in (A) and (G)). 51 nuclei examined. (A) to (J): all experiments at 4 h in meiosis, except *ndt80Δ* (6 h in meiosis).

### Mlh3 associates with meiotic DSB hotspots at a late step of recombination, independently of *NDT80* activation

We next examined Mlh3 association with specific loci by chromatin immunoprecipitation. MutLγ associated with the three tested DSB hotspots during meiosis, reaching a maximum at the expected time of crossover formation (4-5 h) (Figure 3A). Mlh3 also weakly associated with the chromosome axis, where DSB sites transiently relocate during recombination^41,42^ (Figure 3A). MutLγ association with DSB hotspots required DSB formation, since it was not observed in a *spo11Δ* mutant (Figure 3B), but was independent of its nuclease activity (*mlh3-D523N* mutant, Figure 3C). Mlh3 binding also required the ZMM protein, Mer3 (Figure 3D). In *msh4Δ* mutants, Mlh3 recruitment was strongly reduced (Figure 3E), consistent with the reduced number of Mlh3 foci in *msh4Δ* (Figure 2F and 2H). By contrast, Mlh3 bound at hotspots at nearly wild-type levels in *exo1Δ* mutants, implying that MutLγ contact with MutSγ and recombination sites can occur independent of Exo1 (Figure 3F).

**Figure 3:**
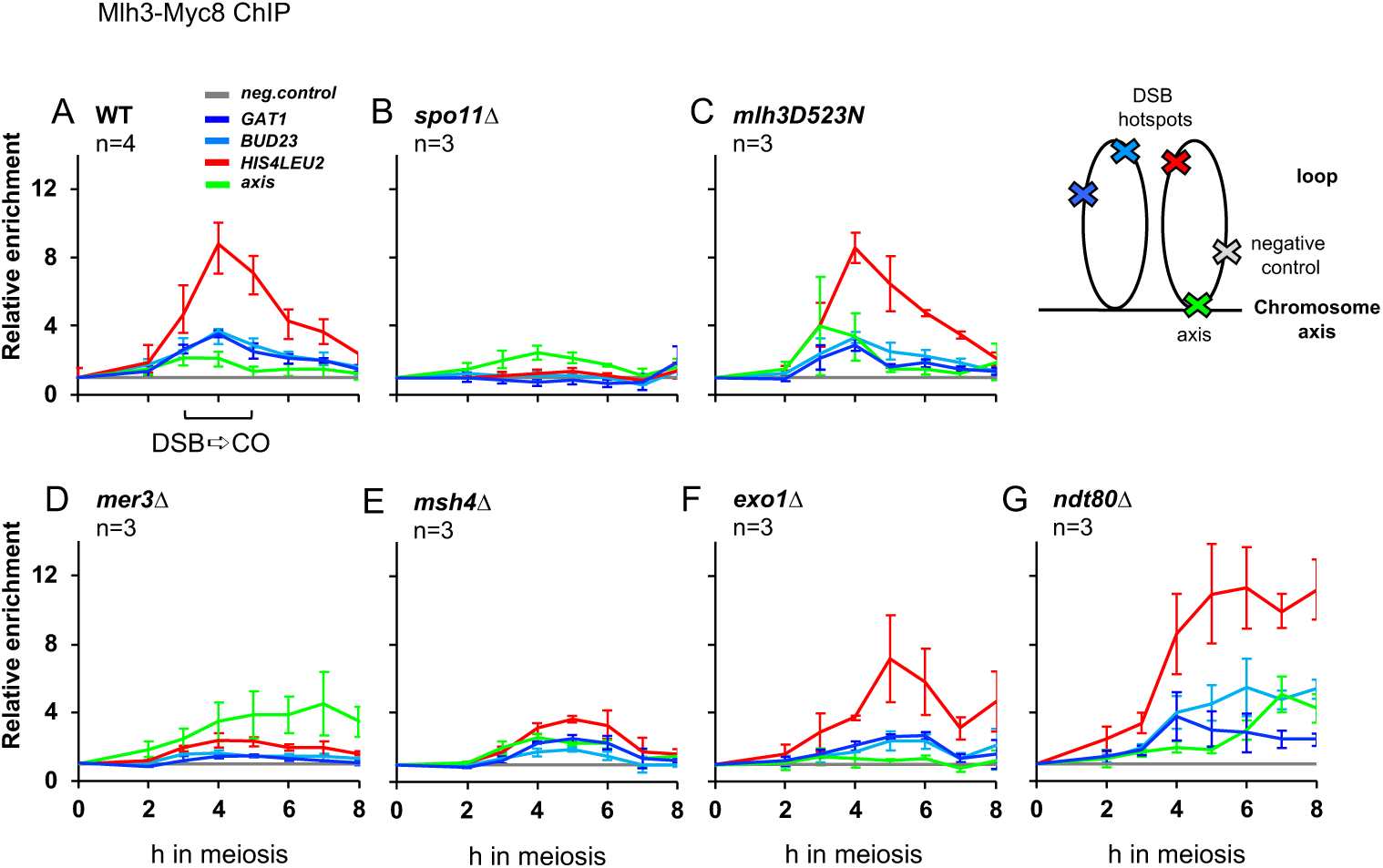
Mlh3 associates with meiotic DSB hotspots at a late step of recombination, independently of *NDT80* activation (A) to (G) Mlh3-Myc levels at the three indicated meiotic DSB hotspots and one axis-associated site relative to a negative control site (*NFT1)* assessed by ChIP and qPCR during meiotic time-courses. Values are the mean ± SEM from three (four in WT) independent experiments. The cartoon illustrates the position of sites analyzed by qPCR relative to the meiotic chromosome structure. (A) WT, (B) *spo11Δ*, (C) *mlh3D523N*, (D) *mer3Δ*, (E) *msh4Δ*, (F) *exo1Δ*. (G) *ndt80Δ* strain (VBD1730).

We next assessed the role of Cdc5 in recruiting Mlh3 to recombination sites, by examining the *ndt80Δ* mutant where Cdc5 is not expressed. Mlh3 was recruited and persisted at DSB sites (Figure 3G) and also formed foci on pachytene chromosomes in *ndt80Δ* (Figure 2G). Mlh3 focus numbers were not greater than in wild type, suggesting that preventing MutLγ activation does not affect its recruitment or its turnover (Figure 2J). We conclude that the lack of dHJ resolution in the absence of Cdc5/PLK1 is not due to a failure to recruit Mlh1-Mlh3 to recombination sites, but rather a failure to activate MutLγ once recruited.

### Cdc5 kinase interacts with both MutLγ and Exo1 bound to recombination sites

The relevant targets of Cdc5 that activate crossover formation by the Mlh1-Mlh3 pathway are not known^31^. To address this issue, we asked if Cdc5 interacts with MutLγ and Exo1 during meiotic recombination, using a copper-inducible *pCUP1-* IME1 construct to produce highly synchronized meiotic cultures^43^. At the time when crossovers start to appear (5.5 h, Figure S2), Cdc5 robustly immunoprecipitated Mlh1, Mlh3 and Exo1 (Figure 4A), consistent with a direct activation of MutLγ cleavage by Cdc5. Indeed, Cdc5 associated with recombination sites during meiosis, at the same time as Mlh3 (Figure 4B). Since MutLγ associates with recombination sites in *ndt80Δ* mutants, where Cdc5 is absent (Figure 3G), our data suggest that MutLγ is activated by Cdc5 once it is at recombining sites. To know which of the MutLγ-Exo1 components is a direct target of Cdc5, we assessed their interdependency for *in vivo* coimmunoprecipitation with Cdc5. Strikingly, Cdc5 still associated at normal levels with Mlh1 and Mlh3 in the absence of Exo1, suggesting that MutLγ itself or another of its interacting partners may be a phosphorylation target of Cdc5 (Figure 4C). Moreover, Exo1 still coimmunoprecipitated with Cdc5 in the absence of interaction with MutLγ (in the *mlh1E682A* mutant), demonstrating that Cdc5 also interacts with Exo1 (Figure 4D).

**Figure 4:**
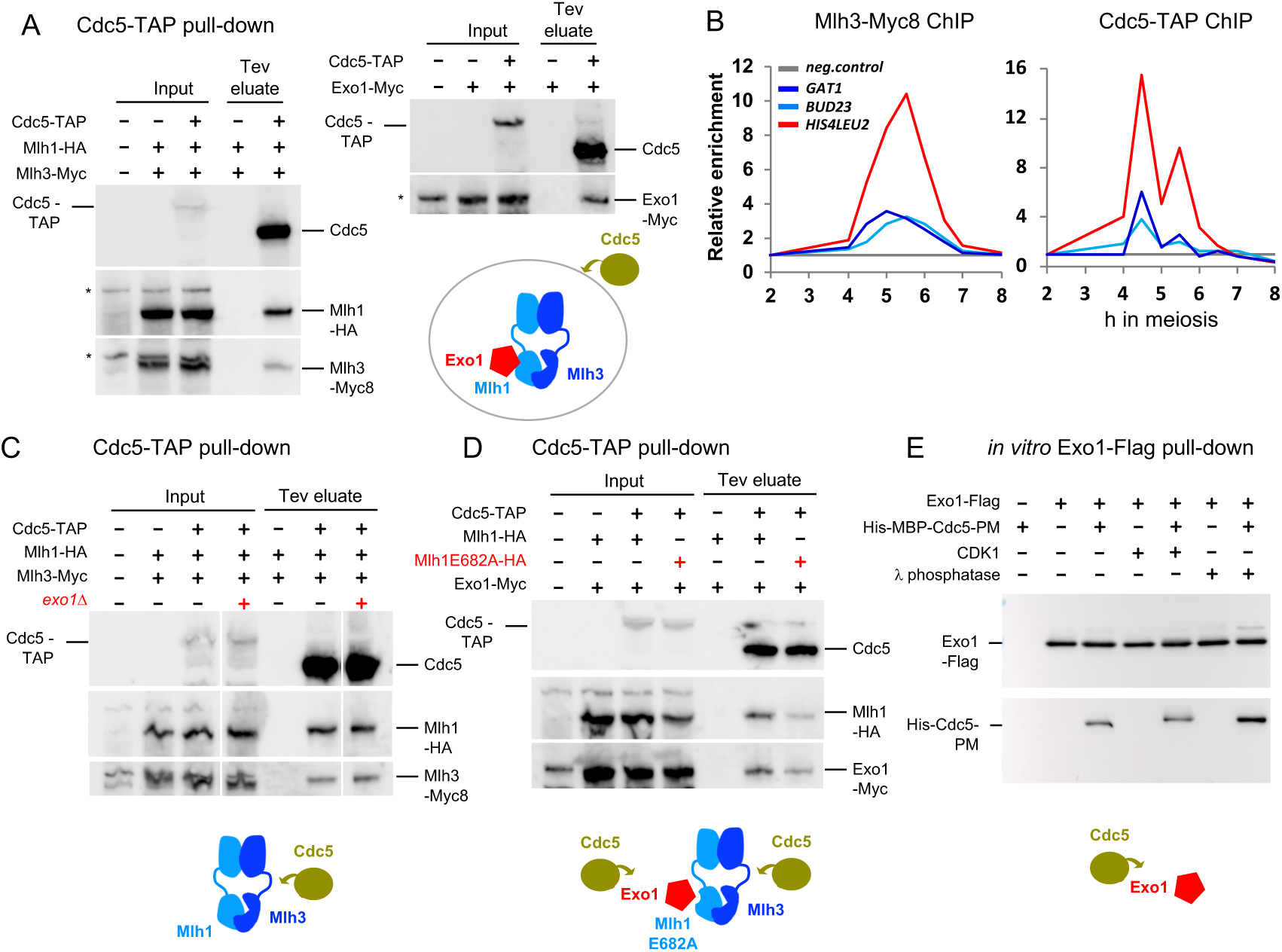
The Cdc5 kinase interacts with both MutLγ and Exo1 bound on recombination sites (A) Coimmunoprecipitation by Cdc5-TAP of Mlh1-HA, Mlh3-Myc and Exo1-Myc from p*CUP1*-*IME1* synchronized cells at 5 h 30 in meiosis analyzed by Western blot. (B) Mlh3-Myc and Cdc5-TAP association with the indicated DSB hotspots as revealed by ChIP and qPCR at the indicated times of a p*CUP1-IME1* synchronized meiotic time-course. Same sites and normalization as in Figure 3. (C) Coimmunoprecipitation by Cdc5-TAP of Mlh1-HA and Mlh3-Myc is independent of Exo1. Same conditions as in (A). (D) Coimmunoprecipitation by Cdc5-TAP of Exo1 is independent of Exo1 interaction with MutLγ. Same conditions as in (A). (E) Direct, phosphorylation-independent interaction between recombinant Exo1 and Cdc5 proteins. Western Blot showing the pulldown of purified Cdc5-His by Exo1-Flag, in the presence or absence of CDK1 or lambda phosphatase. See also Figures S2 and S3.

### Cdc5 directly interacts with Exo1 to promote crossover formation

Cdc5 usually recognizes its substrates through its polo-box domain that preferentially binds a priming phosphate, often formed by CDK1, in a consensus motif of S-S(Phos)/T(Phos)-P^44–46^. Neither Mlh1 nor Mlh3 contain this consensus motif, but Exo1 contains one (SSP residues 663-665), so we focused on Cdc5-Exo1 interaction. We assessed *in vivo* Exo1 phosphorylation by mass spectrometry of Exo1-TAP purified from meiotic cells after induction of Cdc5, and identified S664 as being phosphorylated, along with eight other sites, including three previously described as resulting from DNA damage checkpoint activation^47^(Figures S3A and B), raising the possibility that Cdc5 recognizes this CDK-phosphorylated site. However, mutation of this residue had no effect on crossovers (Figure S3C), suggesting that Cdc5 might interact with Exo1 in a way that does not use this priming phosphosite. This was the case, since purified Exo1 strongly interacted with purified Cdc5, regardless of Exo1 phosphorylation status (Figure 4E). A few other cases of non-canonical, phosphorylation-independent Cdc5 interactions have been described, including the interaction between Cdc5 and Dbf4^48–50^.

We next examined how Cdc5 and Exo1 interact, using yeast two hybrid assays. The polo-box domain of Cdc5 was sufficient for interaction with Exo1, as was found for Dbf4^50^ (Figure 5A). Domain mapping defined a small region of Exo1 (531 to 591) necessary and sufficient for interaction with Cdc5 (Figure 5A). This region contains a motif (570-578) conserved among yeast species, resembling the “RIGA” motif found in Dbf4 (Figure 5B)^50^. Furthermore, mutation of the key residues of this motif (*R570E I572D G574A*, hereafter called *exo1-cid*, Cdc5 interaction-deficient) abolished the two-hybrid interaction between Exo1 and Cdc5 (Figure 5A and S4). Interaction between Exo1-cid and Cdc5 was also reduced in *in vivo* coimmunoprecipitation assays (Figure 5C).

**Figure 5:**
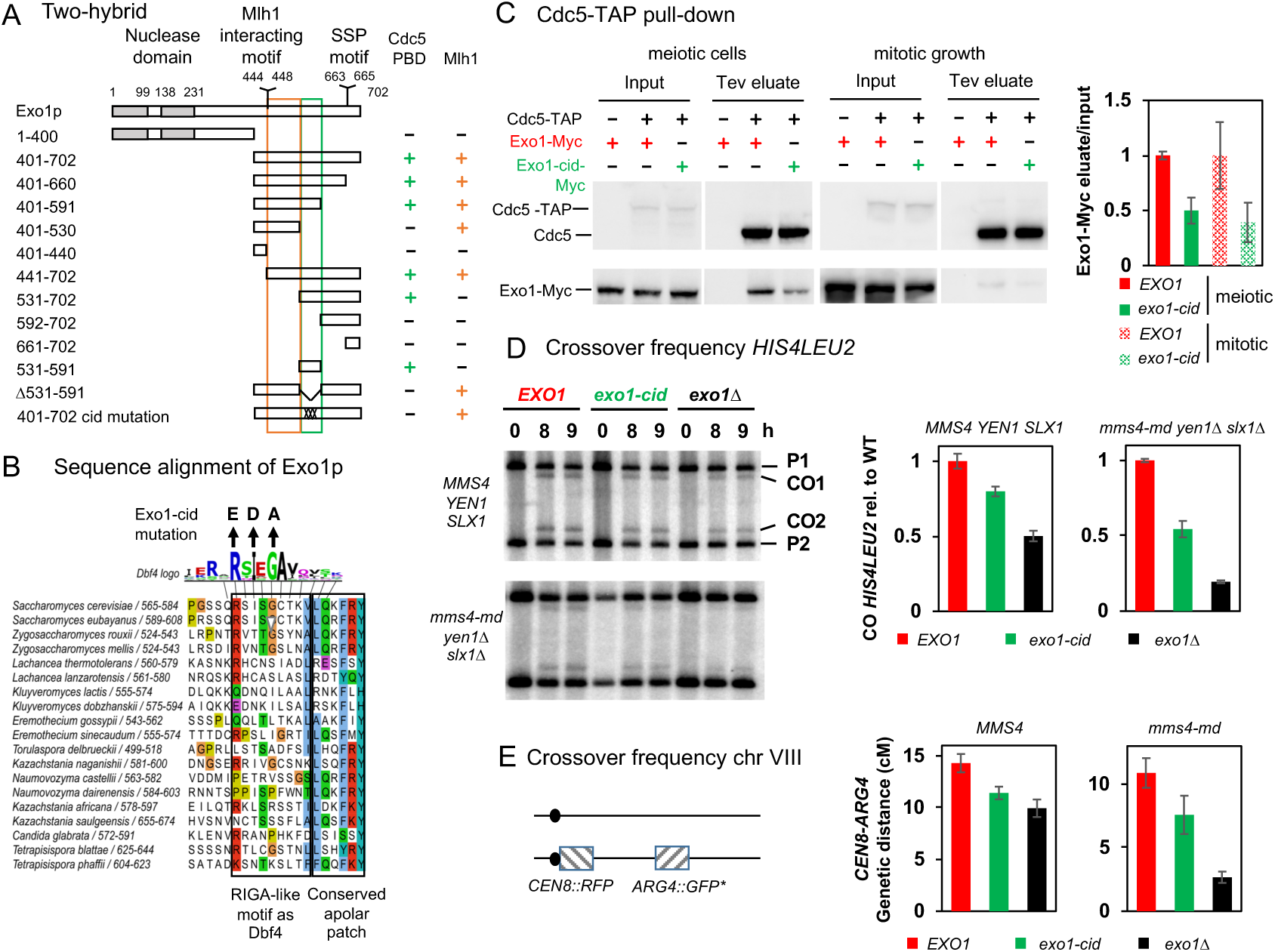
Cdc5 directly interacts with Exo1 to promote crossover formation (A) Delineation of the Exo1 motif responsible for interaction with Cdc5 polo-box domain (PBD) by two hybrid assays. The GAL4-BD fusions with indicated Exo1 fragments were tested in combination with a GAL4-AD-Cdc5-PBD fusion. + indicates an interaction. Exo1-cid: <u>C</u>dc5 interaction-<u>d</u>eficient. (B) Conservation of the Exo1 region interacting with Cdc5 PBD and illustration of the Exo1-cid mutation. The Dbf4 motif interacting with Cdc5 PBD^50^ is indicated (consensus from the 17 Saccharomycetaceae family species). (C) Coimmunoprecipitation by Cdc5-TAP of Exo1-Myc in meiotic cells or in cells growing mitotically. Same conditions as in Fig. 4A. Right: quantification of Exo1-Myc levels in the Tev eluate relative to the input. Error bars represent S.D. of two technical replicates. (D) Meiotic crossover frequencies at the *HIS4LEU2* hotspot in the *exo1-cid* mutant. Left: representative Southern blot analysis of crossovers in the indicated *exo1* mutants, in an otherwise wild-type (*MMS4 YEN1 SLX1)* or triple nuclease mutant (*mms4-md yen1Δ slx1Δ*) background. *mms4-md* stands for *pCLB2-mms4*. Right: quantification of crossovers. Values are the mean±S.E.M of 4 independent experiments (WT background) or mean±S.D.of 2 independent experiments (triple nuclease mutant), normalized to the corresponding *EXO1* value. (E) Meiotic crossovers on chromosome VIII. Left: illustration of the fluorescent spore setup^52^. Right: genetic distances measured in the *CEN8-ARG4* genetic interval, for each indicated genotype. See also Figure S4.

Importantly, *exo1-cid* mutants showed decreased crossovers, specifically those promoted by MutLγ, on two different tested chromosomes, at the *HIS4LEU2* hotspot on chromosome III (Figure 5D), and in the *CEN8-ARG4* interval on chromosome VIII (Figure 5E). Crossover levels in *exo1-cid* were greater than in the *exo1Δ* mutant, but decreased to the same extent as in mutants that impair Exo1 interaction with MutLγ (*exo1-FF477AA* and *mlh1E682A* mutants^27^), highlighting that the *in vivo* crossover function of Exo1 is lost in the *exo1-cid* mutant.

Mutation of all nine phosphorylated residues of Exo1 (mutant *exo1-9A*) did not affect crossovers (Figure S3C). Finally, no phosphoshift of Exo1 was detected at the time of Cdc5 expression (our results and^51^). These results combine to suggest that Cdc5 does not phosphorylate Exo1 itself, or if it does, it is not important for crossover formation. This suggests that Exo1 acts as a bridge between Cdc5 and MutLγ, to recruit Cdc5 to recombination sites and to activate crossover formation.

### Mlh3 distribution on recombination sites is influenced by the length of resection tracts

Mlh3 immunoprecipitates also contained Pms1, the partner of Mlh1 in the MutLα heterodimer, and we confirmed this interaction in reverse Pms1 pulldowns (Figure 1B, Table S1 and Figure S5A). This was unexpected, because two-hybrid data indicate that Mlh3 does not interact with Pms1^23^. However, this coimmunoprecipitation might be mediated by interactions between the Mlh1-Pms1 and Mlh1-Mlh3 heterodimers (Figure S5B). This possible MutLα-MutLγ interaction prompted us to ask if MutLγ-MutLγ interactions also occur, possibly as a result of the proposed MutLγ polymerization needed to cleave DNA *in vitro*^24^. However, we failed to detect any Mlh3-Mlh3 interaction in coimmunoprecipitation assays, suggesting either that MutLγ does not interact with itself *in vivo* or that this may be regulated and occur only at the time of CO formation, making it difficult to detect (Figure S5C).

As a complementary approach, we asked if the shape of Mlh3 distribution around DSB sites is compatible with MutLγ polymerization. For this, we mapped Mlh3 binding sites by ChIPseq, using highly synchronized p*CUP1-IME1* meiotic cultures at the 5 and 5.5 h time-points, when Mlh3 binding at recombination sites reaches a maximum (Figure 4B). We also mapped DSB sites by sequencing Spo11 oligonucleotides from cultures synchronized in the same manner, at 5 h when DSB levels are maximal (Figure S2). The Mlh3 binding map was highly similar to thos of the Mer3 and Zip4 ZMMs (correlation coefficients 0.8) (Figures 6A and S6A). Like Mer3 and Zip4, Mlh3 formed peaks around DSB hotspots, and like Mer3, Mlh3 was weakly associated with axis binding sites (Figures 6A and 6B). Globally, the Mlh3 distribution was dominated by binding to DSB hotspots, and was highly correlated with DSB sites and not with axis sites (Figure S6A). We identified 1155 Mlh3 peaks (Table S2) that we compared to DSB hotspots (Table S3). 484 Mlh3 peaks overlapped at least one of the strongest 1000 DSB hotspots. At the strongest 500 DSB hotspots (mean width 0.5 kb), Mlh3 peaks had a mean width of 2.8 kb, similar to that of Zip4 (2.9 kb) (Figure 6D). The broader distribution of Mlh3 within peaks, compared to that of Spo11 cleavage sites, is consistent with the expected size of recombination intermediates. One determinant of the width of recombination intermediates is the extent of DSB end resection. Consistent with this, the mean width of all Mlh3 peaks overlapping a DSB hotspot (2.2 kb) was similar to that previously determined for the length of resection tracts at DSB hotspots (1.6 kb)^53^ (Figure 6C). To ask if recombination intermediates width determines the extent of Mlh3 binding, we mapped Mlh3 in an *exo1-D173A* (*exo1-nd*) catalytic mutant that shows a 2.2 fold reduction in resection tracts lengths^27,53^. In this mutant, MutLγ peaks at DSB hotspots were clearly less wide than in wild-type (Figure 6C and 6D). The mean width of Mlh3 peaks at all DSB hotspots was 1.8 kb (1.2 fold reduction), and was 2.1 kb (1.4 fold reduction) at the strongest 500 DSB hotspots (Figure 6D).

**Figure 6:**
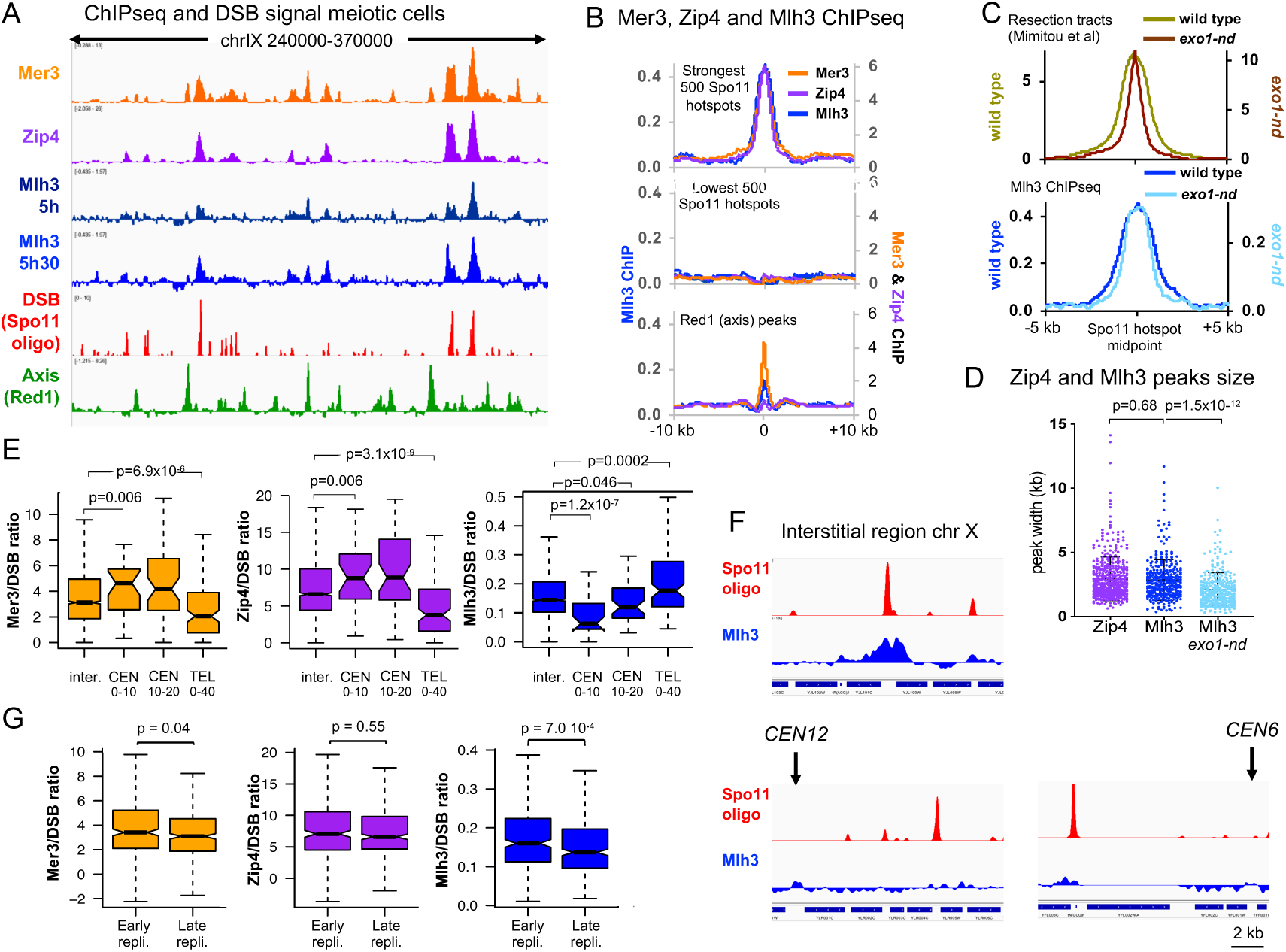
Mlh3 distribution on recombination sites is influenced locally by the length of resection tracts and globally by specific chromosome features (A) ChIPseq binding of Mlh3 at two close time-points compared to the binding of the Mer3 and Zip4^60^ ZMMs, to DSBs (Spo11 oligos,^61^) and to axis sites (Red1 ChIPseq,^62^). Normalized data are smoothed with a 200 bp window. (B) Average ChIPseq signal at the indicated features. Same data as in (A). The Mer3 and Zip4 ChIPseq signals are aligned on the Spo11 hotspots midpoints from^61^, and Mlh3 ChIPseq signal on the p*CUP1-IME1* Spo11 hotspots midpoints (this study). Lower panel, ChIPseq signal is aligned on the Red1 peaks summits from^62^. (C) Average ChIPseq signal of Mlh3 at the 5 h 30 timepoint compared to the distribution of resection tracts computed from resection endpoints (data from^53^, see Extended Methods). Resection tracts are measured at the 500 strongest Spo11 hotspots from^61^, whereas Mlh3 ChIPseq signal (smoothed with a 200 bp window) is aligned on the 500 strongest p*CUP1-IME1* Spo11 hotspots (this study). (D) Width of Zip4 and Mlh3 peaks at the DSB hotspots. Only peaks that match with at least one of the strongest 500 Spo11 hotspots are considered (for Zip4, 421 peaks (Spo11 hotspots from^61^; for Mlh3 and Mlh3 *exo1-nd*, 318 and 300 peaks, respectively (p*CUP1-IME1* Spo11 spots)). A Mann-Whitney-Wilcoxon test was used to compare the datasets. (E) ZMM and Mlh3 signals per DSB vary with the proximity to a centromere or a telomere. The ChIPseq signal of each protein divided by the corresponding Spo11 oligo signal (Spo11 signal from^61^ for Mer3 and Zip4, and p*CUP1-IME1* Spo11 oligo for Mlh3) was computed on the width plus 1 kb on each side of the strongest 2000 hotspots^61^ at the indicated chromosome regions: inter (further than 10 kb from a centromere or 20 kb from a telomere, 1955 hotspots); CEN 0-10kb (29 hotspots); CEN 10-20kb (50 hotspots); TEL 0-40 kb (117 hotspots). Data are represented as boxplots, and the statistical differences (Mann-Whitney-Wilcoxon test) between different regions are indicated. (F) Examples of DSB (p*CUP1-IME1* Spo11 oligo) and Mlh3 binding in an interstitial region and two pericentromeric regions. The normalized signal smoothed with a 200 bp window size is indicated. The scale for Spo11 oligos as well as that for Mlh3 signal are the same in the four graphs to allow a quantitative comparison between the four regions. (G) The ZMM and Mlh3 signals per DSB vary with the timing of DNA replication. Same legend as in (E) for the indicated chromosomal regions. See also Figure S5.

Altogether, our results are consistent with the detectable MutLγ distribution *in vivo* not being much wider than the limits of recombination intermediates and therefore not showing evidence of extensive polymerization over several kilobases. If this occurs, as suggested by activity tests *in vitro*, it may be highly transient and triggered at the precise time of resolution^24,51^.

### MutLγ binding levels correlate well with DSB frequencies, but are reduced in centromeric and late replicating regions

Whether all types of chromosomal regions are equally prone to MutLγ-dependent crossover is an important question. To address this, we used our genome-wide maps of Mlh3 binding and of DSBs to compare the steps of recombination initiation with crossover resolution. Previous studies have shown that small chromosomes tend bind more DSB-forming proteins, and consequently form more DSBs and more crossovers per kb than longer chromosomes^41,54–59^. Consistent with this, Mlh3 density increased as chromosome size decreased (Figure S6B). Globally, the frequency of DSB at a site was highly predictive of a Mlh3 binding. Among the strongest 200 DSB hotspots, 164 had a detected Mlh3 peak (Figure S6A and S6C), and DSB and Mlh3 peaks ranks were significantly positively correlated (r = 0.50, p < 2.2e-16). However, we observed a significant reduction of Mlh3 binding in relation to DSB signal in pericentromeric regions relative to interstitial regions (Figures 6E and 6F and Figure S7A) that was not seen for the two upstream ZMM proteins, Mer3 and Zip4 (Figure 6E). By contrast, in subtelomeric regions, the Mlh3/DSB ratio was not reduced compared to interstitial regions, but the equivalent ratio was reduced for Mer3 and Zip4 (Figure 6E). Thus, these two different non-interstitial chromosomal regions behave differently from most of the genome, but likely for different underlying reasons (see Discussion).

We also asked if the binding of Mer3, Zip4 and Mlh3 on DSB hotspots is modulated by replication timing (see Materials and Methods). Mer3/DSB ratios were slightly lower in late compared to early replicating regions, and Zip4/DSB ratios were similar in both kinds of regions (Figure 6G). By contrast, Mlh3 binding per DSB was reduced in late replicating regions (Figure 6G); this effect was seen in samples from both 5 h (Figure S7B) and 5.5 h (Figure 6G) suggesting that this effect is not due to temporal differences in Mlh3 loading between early and late replicating regions. (Figure S7B).

## Discussion

In this study, we provide the first thorough *in vivo* characterization of the MutLγ complex involved in meiotic crossovers. We show that MutLγ forms foci on meiotic chromosomes and that globally, its binding to recombination sites is repressed close to centromeres and in late replicating regions. This indicates a role of the chromosome structure in modulating the outcome of meiotic recombination. Relevant to MutLγ activation, we show a weak, possibly very transient physical interaction with MutSγ during meiosis and a robust, constitutive interaction with Exo1. We reveal a new non-catalytic function of Exo1 in recruiting Cdc5 to activate crossover formation, which involves a non-canonical, direct interaction, which does not seem to be accompanied by Cdc5-mediated Exo1 phosphorylation. Exo1 serves therefore as a matchmaker between Cdc5 and MutLγ and its partners. To our knowledge, this represents an unprecedented mode of recruiting Cdc5/PLK1 to its phosphorylation targets (Figure 7).

**Figure 7:**
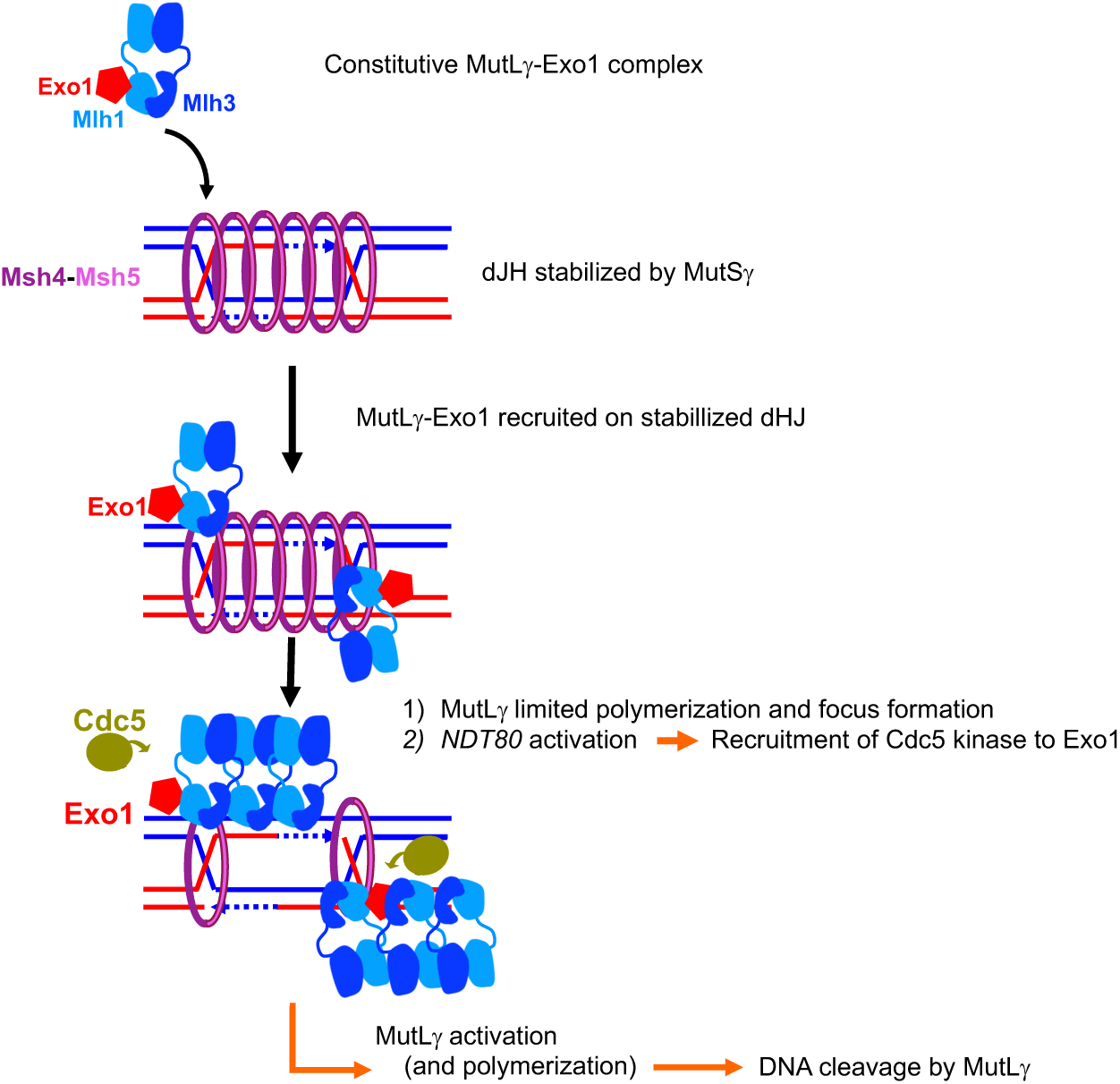
Model of MutLγ binding to sites of recombination and activation for crossover formation MutLγ, in complex with Exo1, binds dJH intermediates that have been stabilized by MutSγ and other ZMM proteins, and may undergo limited polymerization. Upon *NDT80* activation, the Cdc5 kinase is induced, and directly interacts with Exo1. This interaction activates the cleavage of the dHJ substrates by MutLγ and produces crossover formation, which may involve more extensive transient polymerization. The cleavage may consist of a nick, that would result in dJH resolution if another nick is present on the opposite strand, or if the newly synthesized DNA (dotted lines) was not ligated to the parental strand before dHJ resolution.

### The map of Mlh3 binding sites is highly correlated with that of DSBs except in specific chromosome regions

The choice to make a crossover during meiotic DSB repair may be modulated along chromosomes, but a precise and high resolution comparison of DSB and crossover levels is missing (reviewed in^63^). In mice, studies of a few hotspots suggest that crossover/noncrossover ratios can vary from one hotspot to another^64–67^. Recent data comparing a genome-wide map of DSBs with existing genetic maps have shown that in the female mice, crossover formation is repressed at subtelomeric regions^68^. Improper placement of crossovers in the vicinity of centromeres negatively influences meiotic chromosome segregation^69–71^, and crossover formation close to centromeres is infrequent in many species, including humans (reviewed in^72,73^). Although recombination close to centromeres is already reduced at the level of DSB formation, residual DSBs increase the risk of aneuploidy^58^. One way to prevent crossover formation by DSBs that form in pericentromeric regions is to repair them either from the sister chromatid and/or as noncrossovers^54,74^. We found that Mlh3/DSB ratios are lower in pericentromeric regions than along chromosome arms. By contrast, ZMM binding at pericentromeric DSBs is not significantly reduced. We thus propose that at pericentromeres, there is a deselection of ZMM-bound recombination sites, and they are preferentially repaired without MutLγ binding, as noncrossovers. In this regard, the phosphatase PP2A associated to Shugoshin has been shown to counteract cohesin kinases, including Cdc5, in pericentromeric regions^75^. It may also prevent Cdc5-driven MutLγ activation in these regions.

At subtelomeric regions, Mlh3/DSB ratios are not lower than in interstitial regions, but ZMM binding per DSB is reduced. This is counterintuitive since ZMM binding is thought to precede MutLγ binding. One way to interpret these results is that these regions bind ZMM proteins later than interstitial regions and were not detected at their maximum level. This will require further investigation.

Prior to our study, it was not clear if replication timing also influenced the choice to repair DSBs as a crossover^76,77^. We show that, on a genome-wide level, Mlh3 binding per DSB is less frequent in late replicating regions. There may be a preferred time window for loading MutLγ onto recombination intermediates, such that DSBs occurring late as a consequence of late replication would be less likely to load MutLγ. It remains to be determined if these DSBs are repaired by the other nucleases able to generate crossovers (such as Mus81-Mms4) or if they are repaired through the noncrossover pathway. Recent cytological data in tomato suggested that the different crossover promoting nucleases (MutLγ and the structure-specific nucleases) are differently distributed along chromosomes^78^.

### Mlh3 associates with DSB sites, at a late step of recombination, and forms foci distinct from ZMM-bound foci

MutLγ is predicted to act on recombination intermediates that have been stabilized by ZMM proteins, consistent with our finding that, in their absence, Mlh3 binding is absent (*zip3Δ* and *mer3Δ*) or reduced (*msh4Δ*). It is intriguing that we do not see much colocalization between ZMM and Mlh3 foci. Two possibilities emerge: either there are two types of ZMM-crossover sites, those that have Zip3 and Msh4 and those that have MutLγ, or, alternatively, Zip3 and Msh4 foci precede Mlh3 foci on all sites, but overlap only for a short time after which Zip3 and Msh4 disappear. Since the association of Mlh3 with chromosomes and recombination sites depends on ZMMs, we propose that ZMMs may preserve the dHJs and hand them over to MutLγ, thereby positioning and activating MutLγ on the recombination intermediates, without staying on chromatin as visible foci when the Mlh3 focus forms. Our results are consistent with those in mice, where early co-localization between Msh4 and Mlh1 foci disappears at mid-pachytene (Kneitz et al., 2000; Santucci-Darmanin et al., 2002).

Another surprising feature is that the average number of Mlh3 foci (16) detected is lower than for its precursors, Zip3 or Msh4 (around 45). All studies including this one have converged to show that ZMM foci numbers in budding yeast are lower than those of DSBs (around 200 per meiosis), and since genetic data predict around 75 ZMM-dependent crossovers per meiosis^57,63^, it was proposed that all ZMM foci give rise to crossovers, contrary to plants, mammals and *Sordaria*^62,79–81^. We envisage two possibilities to explain this result. Either a similar selection of a subset of ZMM-bound sites operates in budding yeast, to become bound by Mlh3. ZMM-foci remaining on the intermediates that have not become bound by MutLγ may be dismantled by helicases and give rise to noncrossover products. In *Sordaria*, Msh4 foci numbers diminish between early and mid-pachytene, a time frame in which Mlh1-Mlh3 is believed to be recruited (Storlazzi et al., 2010). Another possibility is that Mlh3 residence time on late crossover intermediates may be limited and therefore Mlh3 is not detected as a focus within a single nucleus on all crossover sites.

### MutLγ forms a constitutive complex with Exo1 in meiotic cells but interacts only transiently with MutSγ

Our finding that MutLγ is constitutively associated with Exo1 is reminiscent of MutLα function in mismatch repair, and indeed the same site on Exo1 is used for interaction with MutLα and with MutLγ, even though Exo1 has non-catalytic function in meiotic crossover formation (this study and^26,27^). The relationships with Msh4-Msh5 are less clear. In many organisms (budding yeast, mice and plants) *msh4* or *msh5* mutants have an earlier meiotic defect than MutLγ mutants, but it is attractive to propose, by analogy to mismatch repair, that MutSγ works also directly with MutLγ-Exo1 for crossover formation, and thus that MutSγ should have both early and late functions. Consistent with an early role for MutSγ, we show that most Msh4 foci do not colocalize with Mlh3. It is worth noting that, in *C. elegans*, MutSγ promotes crossovers although crossover resolution is not achieved by MutLγ, suggesting that, in this organism, either MutSγ performs only an early crossover formation function, or that it cooperates with a nuclease other than MutLγ^82,83^. On the other hand, the interaction between MutLγ and MutSγ in meiotic cells, which we detected by coimmunoprecipitation, fits with a late function for MutSγ through direct activation of MutLγ, although the fraction of Msh4 interacting with Mlh3 is quite small. A similar interaction was also described for the cognate mouse proteins^84,85^. At this time, it is difficult to determine if MutSγ contributes mainly by creating the proper substrate for an asymmetric dHJ resolution by MutLγ, or if it also directly activates MutLγ catalysis (Figure 7).

### Does MutLγ polymerize on recombination intermediates *in vivo*?

We did not detect any MutLγ self-interaction. In addition, our data indicate that the distribution of Mlh3 results mostly from a passive process that adopts the size of recombination intermediates. This seems incompatible with extensive polymerization over several kilobases, which would be a very active process, but is compatible with a more limited occupancy by several MutLγ complexes (Figure 7). Recently, it was shown that during meiotic crossover formation, two MSH-5 foci are present at crossover sites as doublets, consistent with two sets of MutSγ complexes accumulating on each DNA duplex in the interval between the two Holliday junctions^82^. These may be the sites MutLγ acts on, and would fit well with the observed distribution of MutLγ on recombination sites. We cannot exclude that once loaded, MutLγ may polymerize transiently when becoming active on a region broader than recombination intermediates (Figure 7). It has recently been proposed that the Chd1 chromatin remodeler Chd1 may promote such a mechanism^51^. Further experiments examining Mlh3 dynamics at the time of dHJ resolution will be required to examine this possibility.

### Exo1 serves as a Cdc5 recruiting platform for MutLγ crossover formation

Our findings have important insights for the step of crossover formation. First, we show that MutLγ binding at recombination sites does not require Exo1. This is similar to the situation in mice, where Mlh1 foci are still observed in the absence of Exo1^30^. Second, since Exo1 is constitutively associated with MutLγ, it is unlikely that the binding of Exo1 to MutLγ activates resolution. Rather, our data, in particular the observation that mutants blocking Exo1-Cdc5 interaction have crossover defects, suggest that resolution is activated by Cdc5 recruitment to MutLγ-bound intermediates, through a direct interaction with Exo1.

One way this could happen would be direct Cdc5 phosphorylation of its interacting partner, Exo1. However, this seems unlikely. We did not detect a change in Exo1 electrophoretic mobiliy as a result of Cdc5 activation, and mutation of Exo1’s known phosphorylated residues had no effect on crossovers. There are cases where Cdc5 directly binds to one protein in a complex but phosphorylates another target, but this usually also involves the phosphorylation of the Cdc5 binding partner^86^. To our knowledge, our study is the first example of the binding partner of Cdc5 not being phosphorylated. In addition, we show that the interaction of Cdc5 with Exo1 is phosphosite-independent, and involves the same kind of motif as the phosphorylation-independent interaction of Cdc5 with Dbf4. This opens new avenues to search for other such partners of Cdc5 that may contain similar interacting motifs, as well as for partners of other polo-like kinases, which are highly conserved between organisms. Strikingly, target recognition (for Dbf4 and Exo1) involves the same Cdc5 region (the PBD) that recognizes phosphopeptides. Structural information would be useful to decipher if these two modes involve different binding interfaces in the Cdc5 PBD.

Cdc5 function in MutLγ crossovers involves phosphorylation of at least a substrate, because induction of a kinase-dead Cdc5 in meiosis is not sufficient to promote crossover formation^31^. An obvious candidate for phosphorylation, once Cdc5 is recruited by Exo1, is the MutLγ complex itself. However, attempts failed to detect any Cdc5 activation-associated change in the mobility of Mlh1 or Mlh3 due to (this study and^51^), and published meiotic phosphoproteomes datasets do not contain meiotic phosphorylation sites on Mlh1 or Mlh3 that would appear upon Cdc5 induction^87^. Another possible target is the Chd1 protein, recently described as a binding partner of MutLγ-Exo1 and important for MutLγ crossovers^51^. Alternatively, or in addition, it is possible that binding of Exo1 to Cdc5 may cause a conformational change that activates MutLγ for crossover formation.

As a whole, our study reveals that MutLγ activation is tightly controlled *in vivo*, both locally through the direct coupling of crossover formation with key meiotic progression steps by the Cdc5 kinase, and globally by the underlying chromosome structure. Much of this regulation is likely to operate in mammals, where the key proteins are conserved. Our study provides an illustration of how nucleases at risk of impairing genome integrity in the germline are kept in check through multiple control levels.

## Methods

### Yeast manipulations

All yeast strains are derivatives of the SK1 background are listed in Table S4, and their place of appearance in the figures is in Table S5. All experiments were performed at 30°C. Two different approaches were used for meiosis induction. In the first one, cells were grown in SPS presporulation medium and transferred in sporulation medium as described^88^. For highly synchronous copper inducible meiosis, the procedure was described in^43^. Cells were grown in YPD to exponential phase and were inoculated at a starting 0.05 OD into reduced glucose YPD (1% yeast extract, 2% peptone, 1% glucose) and grown to an OD_600_ = 11.0-12.0 for 16-18 h. Cells were washed, resuspended in sporulation medium (1.0% (w/v) potassium acetate, 0.02% (w/v) raffinose, 0.001% polypropylene glycol) at OD_600_ of 2.5. After 2 h, copper (II) sulfate (50 µM) was added to induce *IME1* expression from the *CUP1* promoter. For strain constructions and spore viability measurements, sporulation was performed on solid sporulation medium for two days.

### Yeast strains construction

Yeast strains were obtained by direct transformation or crossing to obtain the desired genotype. Site directed mutagenesis, internal and C-terminal tag insertions and gene deletions were introduced by PCR. All transformants were confirmed using PCR discriminating between correct and incorrect integrations and sequencing for epitope tag insertion or mutagenesis. For Mlh3 internal tagging, the Myc8, Myc18 or His6-Flag3 (referred to as Flag) tag sequence, flanked on each side by a GGGGSGGGGS linker sequence, was inserted between residues 512 and 515 of Mlh3 (513 and 514 were deleted). For Pms1 internal tagging, the His6-Flag3 tag sequence, flanked on each side by a GGGGSGGGGS linker sequence, was inserted between residues 758 and 761 of Pms1 (759 and 760 were deleted). Cdc5, Exo1 and Msh4 were C-terminally tagged with the TAP tag sequence^89^. Exo1 was tagged with 13 copies of the Myc epitope and Msh4 with 3 copies of the HA epitope using a PCR-based strategy^90^. The KanMX-*pCUP1-3HA-IME1* locus was PCR-amplified from 33604 strain from A. Amon lab and used for direct transformation. The *exo1-D173A* (*exo1-nd*) mutation was introduced by CRISPR-Cas9 mediated cleavage, as described^40^.

### Analysis of crossover frequencies

For crossover analysis at *HIS4LEU2*, cells were harvested from meiotic time courses at the indicated time point. 2 μg genomic DNA was digested with *Xho*I and analyzed by Southern blot using a labeled DNA probe A as described^6^. The radioactive signal was detected by a Phosphorimager (Typhoon, GE Healthcare) and quantified using the Image Quant software as described^40^. For genetic distances on chromosome VIII, diploids were sporulated in liquid medium, and recombination between fluorescent markers was scored after 24h in sporulation, by microscopy analysis as described^52^. Two independent sets of each strain were combined and at least 600 tetrads were counted. Genetic distances in the *CEN8-ARG4* interval were calculated from the distribution of parental ditype (PD), nonparental ditype (NPD) and tetratype (T) tetrads and genetic distances (cM) were calculated using the Perkins equation: cM = (100 (6NPD + T))/(2(PD + NPD + T)). Standard errors of genetic distances were calculated using Stahl Lab Online Tools (https://elizabethhousworth.com/StahlLabOnlineTools/).

### Flag-affinity pull-down and mass spectrometry analysis

2.10^10^ cells at 4 h in meiosis were harvested, washed two times with ice-cold TNG buffer (50 mM Tris/HCl pH 8; 150 mM NaCl, 10% Glycerol; 1 mM PMSF; 1X Complete Mini EDTA-Free (Roche); 1X PhosSTOP (Roche)) and flash-frozen in liquid nitrogen. Frozen cells were mechanically ground in liquid nitrogen with the 6775 Freezer/Mill cryogenic grinder (SPEX SamplePrep). The resulting powder was resuspended in 25 mL of lysis buffer (50 mM Tris/HCl pH 7.5; 1 mM EDTA; 0.5% NP40; 10% glycerol; 150 mM NaCl; 1X Complete Mini EDTA-Free (Roche); 1X PhosSTOP (Roche), 0 or 210 U/mL benzonase (Sigma)) and incubated 1 h at 4°C with rotation. The lysate was cleared by centrifugation at 8000 g for 10 min and pre-cleared by pre-incubated with 100 μl Mouse IgG−Agarose beads (A0919 Sigma) without antibody for 2 h at 4°C on wheel. The pre-cleared lysate was incubated with 100 μl of washed and buffer equilibrated anti-Flag magnetic beads (Sigma-Aldrich, St. Louis, MO) for 2 h at 4°C. The beads were washed once with lysis buffer and three times with washing buffer (20 mM Tris/HCl pH 7.5; 0.5 mM EDTA; 0.1% tween; 10% glycerol; 150 mM NaCl; 5 mM MgCl2; 0.5 mM PMSF; 1X Complete Mini EDTA-Free (Roche, Switzerland); 1X PhosSTOP (Roche)). Proteins were eluted with 5 bed volume of elution buffer (20 mM Tris/HCl pH 8; 0.5 mM EDTA; 0.1% tween; 10% glycerol; 150 mM NaCl; 5 mM MgCl2; 0.5 mM PMSF; 1X Complete Mini EDTA-Free (Roche); 1X PhosSTOP (Roche); 100 μg/mL Flag peptide) for 2 h at 4°C. Proteins were separated by SDS-PAGE, stained with colloidal blue, and bands covering the entire lane were excised for each sample. In-gel digestion was performed overnight by using trypsin/LysC (Promega, Madison, WI). Peptides extracted from each band were analyzed by nanoLC-MS/MS using an Ultimate 3000 system (Dionex, Thermo Scientific, Waltham, MA) coupled to a TripleTOF^TM^ 6600 mass spectrometer (ABSciex). For identification, data were converted to mgf files, merged with Proteome Discoverer (version 2.2) and searched against the Swissprot fasta database containing *S. cerevisiae* (2019_04, 7905 sequences) using Mascot^TM^ (version 2.5.1) and further analyzed in myProMS v3.6^91^. Data are available via ProteomeXchange with the identifier PXD014180^92^. Only proteins found in four experiments and not in the control IPs were considered candidates.

### Coimmunoprecipitation and Western blot analysis

1.2.10^9^ cells were harvested, washed once with PBS, and lyzed in 3 ml lysis buffer (20 mM HEPES/KOH pH7.5; 150 mM (TAP purification) or 300 mM (anti-FLAG immunoprecipitation) NaCl; 0.5% Triton X-100; 10% Glycerol; 1 mM MgCl2; 2 mM EDTA; 1 mM PMSF; 1X Complete Mini EDTA-Free (Roche); 1X PhosSTOP (Roche); 0 or 125 U/mL benzonase) with glass beads three times for 30 s in a Fastprep instrument (MP Biomedicals, Santa Ana, CA). The lysate was incubated 1 h at 4°C. For TAP-tagged proteins precipitation, 100 μL of PanMouse IgG magnetic beads (Thermo Scientific) were washed 1:1 with lysis buffer, preincubated in 100 μg/ml BSA in lysis buffer for 2 h at 4°C and then washed twice with 1:1 lysis buffer. The lysate was cleared by centrifugation at 13,000 g for 5 min and incubated overnight at 4°C with washed PanMouse IgG magnetic beads. The magnetic beads were washed four times with 1 mL of wash buffer (20 mM HEPES/KOH pH7.5; 150 mM NaCl; 0.5% Triton X-100; 5% Glycerol; 1 mM MgCl2; 2 mM EDTA; 1 mM PMSF; 1X Complete Mini EDTA-Free (Roche); 1X Phos-STOP (Roche)). The beads were resuspended in 30 μL of TEV-C buffer (20 mM Tris/HCl pH 8; 0.5 mM EDTA; 150 mM NaCl; 0.1% NP-40; 5% glycerol; 1 mM MgCl2; 1 mM DTT) with 4 μL TEV protease (1 mg/ml) and incubated for 2 h at 23°C under agitation. The eluate was transferred to a new tube. For anti-FLAG immunoprecipitation, 50 μl of Protein G magnetic beads were used and 10 μg of mouse monoclonal anti-FLAG primary antibody M2 (Sigma) were added before overnight incubation at 4°C with the lysate. Wash buffer contained 300 mM NaCL. After washing, beads were resuspended in 25 μl of 2x SDS protein sample buffer. Beads eluate was heated at 95°C for 10 min and loaded on acrylamide gel (4-12% Bis-Tris gel (Invitrogen)) and run in MOPS SDS Running Buffer (Life Technologies). Proteins were then transferred to PVDF membrane using Trans-Blot® Turbo™ Transfer System (Biorad) at 1 A constant, up to 25 V for 45 min. Proteins were detected using c-Myc mouse monoclonal antibody (9E10, Santa Cruz, 1:500), Flag mouse monoclonal antibody (M2, Sigma, 1:1000), HA.11 mouse monoclonal antibody (16B12, Biolegend, 1/750) or TAP rabbit monoclonal antibody (Invitrogen, 1/2000). Signal was detected using the SuperSignal West Pico or Femto Chemiluminescent Substrate (ThermoFisher). Signal was quantified after image acquisition with Chemidoc system (Biorad).

### Recombinant proteins and interaction assays

Recombinant yeast Exo1-Flag purification from Sf9 insect cells was described previously^93^. Mlh1-Mlh3 were prepared as described^20^. The *CDC5* gene was amplified from *S.cerevisiae* genomic DNA (SK1) with added *Nhe*I and *Xho*I sites, and cloned into the *Nhe*I and *Xho*I sites of a vector backbone modified from pFB-MBP-*MLH3*-his^20^ by adding sequence for a 8xHis tag before the MBP gene, to make pFB-His-MBP-*CDC5*. Cdc5 is only active when phosphorylated. We addressed this issue by mutating the conserved threonine 238 (T210 in human PLK) to aspartic acid (T238D)^94^. This mutation is known to activate yeast Cdc5 constitutively^95^. The Cdc5T238D (Cdc5-PM) protein was expressed and purified in Sf9 cells following the protocol described for Mlh1-Mlh3^20^, except that both the His and the MBP tags are present in the recombinant protein. 1 mg of protein was obtained from 1.4 L *Sf*9 insect cells culture (Figure S3D). CDK1-cyclin B was purchased from Merck. Lambda phosphatase was purchased from NEB.

To phosphorylate Exo1-Flag, 1 µg of the recombinant Exo1-Flag was incubated in kinase buffer (50 mM Tris-HCl pH 7.5, 5 mM Mg(OAc)2, 0.2 mM EDTA) supplemented with phosphatase inhibitor mixture (0.5 mM Na3VO4, 10 mM p-nitrophenyl phosphate). 40 ng of CDK1-cyclin B were added to the reaction. Phosphorylation reaction was started by adding 0.1 mM ATP and incubated for 15 min at 30°C. To dephosphorylate Exo1-Flag, 1 µg of the recombinant Exo1-Flag was incubated in 10X NEBuffer for Protein MetalloPhosphatases (PMP) supplemented with 1 mM MnCl_2_. 280 U of Lambda phosphatase were added to the reaction. Dephosphorylation reaction was started by incubating for 15 min at 30°C. To test for the interaction between His-MBP-Cdc5 and Exo1-Flag treated or not with CDK1 or Lambda phosphatase, 10 µl Protein G magnetic beads (Thermo Scientific) were used to capture 4 µg anti-Flag Monoclonal M2 antibody (Sigma, F3165). Exo1-Flag (1µg) was incubated with the beads in 60 µl binding buffer III (25 mM Tris-HCl pH 7.5, 3 mM EDTA, 1 mM DTT, 20 mg/ml BSA, 75 mM NaCl) for 60 min with continuous mixing. Next, the beads were washed 3 times with 150 µl wash buffer III (25 mM Tris-HCl pH 7.5, 3 mM EDTA, 1 mM DTT, 120 mM NaCl, 0.05% Triton X-100). 1 µg His-MBP-Cdc5 was then added to the resuspended beads in 60 µl binding buffer III, and incubated for additional 60 min with continuous mixing. Beads were washed 3 times with 150 ul wash buffer III and boiled for 3 min at 95 °C in SDS buffer to elute the proteins. The protein complexes were detected by western blot with anti-His antibody (Genscript, A00186) and anti-Flag antibody (Sigma, F3165)

### Yeast two-hybrid assays

Yeast two-hybrid assays were performed exactly as described^40^. *EXO1* fragments and *CDC5-PBD* (residues 340 to 705) were PCR-amplified from SK1 genomic DNA. PCR products were cloned in plasmids derived from the 2 hybrid vectors pGADT7 (GAL4-activating domain) and pGBKT7 (GAL4-binding domain) creating N terminal fusions and transformed in yeast haploid strains Y187 and AH109 (Clontech), respectively. Interactions were scored, after mating and diploid selection on dropout medium without leucine and tryptophan, as growth on dropout medium without leucine, tryptophan, adenine and histidine.

### Cytology

4×10^7^ cells were harvested at the indicated time-point and yeast chromosome spreads were prepared as described^96^. Mlh3-myc18 was stained with primary anti-myc rabbit antibody (Abcam, 1:600) and secondary FITC-conjugated anti-rabbit (Thermo Fischer 1:600) for all cytological experiments except for the colocalization with Zip3, for which primary 9E11 anti-myc mouse antibody was used together with a secondary Cy3-conjugated anti-mouse (Jackson ImmunoResearch, 1:600). Msh4-HA3 was stained with primary anti-HA mouse antibody (Anopoli, 1:1000) and secondary Cy3-conjugated anti-mouse antibody (Thermo Fischer 1:600) when used in parallel with rabbit based anti-myc. Zip3 was stained with a primary anti-Zip3 rabbit antibody (1:2000), a gift from Akira Shinohara. The secondary antibody was Cy5-conjugated anti-rabbit (Amersham, 1:500). Fluorescent microscopy was carried out on a ZEISS AXIO Imager M2 with a ZEISS Plan-Neofluar 100x, and a 2x additional magnification by a Zeiss optovar. Images were taken at an exposure of 1 second for DAPI (BFP channel), Cy3 (Cy3 channel), FITC (FITC channel) and 2 seconds for Cy5 (AF660 channel). The Light source: Sola SM II (Llumencor); camera: CoolSNAP HQ2 (Visitron Systems GmbH); acquisition software: Visiview (Visitron Systems GmbH). Nuclei acquired with this setup were analyzed by Fiji software and R-scripts.

### Chromatin immunoprecipitation, real-time quantitative PCR and ChIPseq

For each meiotic time point, 2.10^8^ cells were processed as described^97^, with the following modifications: lysis was performed in Lysis buffer plus 1 mM PMSF, 50 μg/mL Aprotinin and 1X Complete Mini EDTA-Free (Roche), using 0.5 mm zirconium/silica beads (Biospec Products, Bartlesville, OK). We used 1.6 μg of c-Myc monoclonal antibody and 50 μL PanMouse IgG magnetic beads or 1 µl of anti-Flag antibody and 30 µl Protein G magnetic beads (Thermo Scientific). For Cdc5-TAP ChIP, the lysate was directly applied on 50 μL PanMouse IgG magnetic beads. Before use magnetic beads were blocked with 5 μg/μl BSA for 4 h at 4°C.

Quantitative PCR was performed from the immunoprecipitated DNA or the whole cell extract using a 7900HT Fast Real-Time PCR System and SYBR Green PCR master mix (Applied Biosystems, Thermo Scientific) as described^97^. Results were expressed as % of DNA in the total input present in the immunoprecipitated sample and normalized first by the negative control site in the middle of *NFT1*, a 3.5 kb long gene, and then by the 0 h time point of the meiotic time-course or the 2 h time-point for copper-induced synchronous meiosis. Primers for *GAT1*, *BUD23*, *HIS4LEU2*, Axis and *NFT1* have been described^40^. For ChIPseq experiments, 1. 10^9^ cells were processed as described before^60^.

### Spo11 oligonucleotides mapping

Spo11-Flag oligonucleotides were purified and processed for sequencing library preparation as previously described^98^. Briefly, cells were harvested from a 600 ml sporulating culture (VBD2016) after synchronizing in meiosis thanks to the p*CUP1-IME1* system at the 5 h time-point to approximate time of peak Spo11-oligo levels. The experiment was made on two biological replicates. Sequencing (Illumina HiSeq 2500, 2 × 50 bp paired-end reads) was performed in the MSKCC Integrated Genomics Operation. Clipping of library adapters and alignment of reads to S288C using genome version sacCer2 were performed by the Bioinformatics Core Facility at MSKCC using a custom pipeline as described^58,59^. The Spo11 oligo data generated in this study have been deposited at the Gene Expression Omnibus database with accession number GSE133108. The list of hotspots of each replicate is in Table S3. The two duplicates were highly similar (Figure S6) and were pooled for further analyses.

### Illumina sequencing of ChIP DNA and read normalization

Purified DNA was sequenced using an Illumina HiSeq 2500 instrument and following the Illumina TruSeq procedure, generating paired-end 50 bp reads. Mer3-Flag ChIPseq was performed in two independent replicates from cells (strain VBD1421) at the 4 h time-point synchronized using the SPS preculture method^88^. The negative control was from an untagged anti-Flag ChIPseq sample (strain VBD1311). Mlh3-Myc8 ChIPseq was performed once at two different time-points (t = 5 h and t = 5.5 h) on meiotic cells (VBD1837 or VBD2026 for the *exo1-nd* strain) synchronized with the copper-inducible system. The negative control was Mlh3-Myc8 ChIPseq in a DSB-deficient *spo11Δ* strain (VBD1841) at the same respective time-points, to reveal only DSB-associated Mlh3 binding sites. Reads were aligned to the sacCer2 version (SGD June 2008) of the *S. cerevisiae* S288C genome, using Bowtie, allowing for 2 mismatches. Reads that matched more than once in the genome or matching to mitochondrial or ribosomal DNA were eliminated from further analysis. Paired-end extended reads were normalized to the negative control, bigwig files generated and then the negative control was subtracted to generate normalized and background-subtracted bigwig files using a custom script as described^60^.

The ChIPseq data generated in this study have been deposited at the Gene Expression Omnibus database with accession number GSE132850.

### Bioinformatic analyses

Sequencing data were analyzed using https://usegalaxy.org/ and custom bash and R scripts. ChIPseq peak calling was performed on the experiment and negative control Bam alignment files using the MACS2 program at https://usegalaxy.org/, with parameters available upon request. Signal smoothing was performed on the normalized and background-subtracted bigwig files using a custom script employing fast Fourier transform convolution with a sliding window of the size indicated in legends of Figures 6 and S6. The script is available upon request.

For computing the ChIPseq signal per DSB represented for different proteins on Figures 6E, 6G and S7, the ChIPseq signal was multiplied by a factor before subtracting the control signal, to avoid obtaining negative values and negative ratios after dividing by the Spo11 oligo signal, since the ChIPseq signal is very low for some proteins close to centromeres and/or telomeres. This was done for further statistical analyses and does not affect the conclusions. The following factors were used: Mer3: 1.2; Zip4: 1.3; Mlh3 _5 h 30: 1.2; Mlh3 _5 h: 1.15; Mlh3 *exo1-nd*: 1.2. For defining interstitial hotspots in early or late replicating regions, we used the replication index data at the 3.5 h meiotic time-point from^99^, and selected among the strongest 2000 hotspots from^61^ those located further than 20 kb away from a telomere or a centromere, and that had a log2 replication index lower than −0.07 (early regions, 306 hotspots) and those with a replication index higher than 0.07 (late regions, 351 hotspots).

To generate the graph Figure 6C representing the lengths of resection tracts, we used the data from^53^ from wild-type cells at t= 4 h, and deduced the distribution of the tracts from the determination of resection end-points. For signal density and chromosome size, the normalized total Mlh3 ChIPseq or p*CUP1-IME1* Spo11 oligo signal was computed for each chromosome, divided by the chromosome length and plotted as a function of chromosome size. The chrIII was omitted, because the strains used contain the artificial *HIS4LEU2* hotspot that modifies the behavior of chromosome III regarding chromosome size.

## Acknowledgments

We thank Wolf D. Heyer for helpful suggestions, Michael Lichten and Arnaud De Muyt for critical comments on the manuscript, Hajime Murakami for the meiotic replication data, Angelika Amon and Folkert van Werven for the p*CUP1*-*IME1* construct, Hardeep Kaur and Michael Lichten for the p*GAL1*-*CDC5* strain, Rajeev Kumar for constructing the Mlh3-Myc8 allele and Akira Shinohara for the anti-Zip3. We thank the Institut Curie PICT-IBISA Pasteur Imaging facility, member of the France Bioimaging National Infrastructure (ANR-10-INBS-04) and the NGS platform, supported by the grants ANR-10-EQPX-03 and ANR10-INBS-09-08 and by the Cancéropôle Ile-de-France. This work was supported by the Institut Curie and the CNRS; by Agence Nationale de la Recherche (ANR-15-CE11-0011) to V.B., J.-B.C. and P.C.; by the French Infrastructure FRISBI (ANR-10-INBS-05) to J.-B.C, and by “Région Ile-de-France” and the FRM medical grants (to D.L.). A.S. was supported by a postdoctoral fellowship from the Fondation ARC and by the Labex DEEP (ANR-11-LBX-0044).

## Author Contributions

Conceptualization, A.S. and V.B.; Methodology, A.S., C.A., F.R., Y.D., L.R., B.L., D.L., P.C., R.G., F.K., J.B.C. and V.B.; Formal Analysis, A.S., C.A., F.R., Y.D., L.R., B.L., X.M., D.L., R.G., F.K. and V.B.; Investigation, A.S., C.A., F.R., Y.D., L.R., B.L. and X.M.; Writing – Original Draft, A.S. and V.B.; Writing – Review & Editing, A.S., D.L., S.K., F.K., J.B.C. and V.B.; Supervision, S.K, P.C. F.K. and V.B.; Funding Acquisition, V.B. and J.B.C.

## SUPPLEMENTAL TABLES AND FIGURES

**Table S1. Mlh3-Flagged pulled-down proteins (Excel spreadsheet)**

**Table S2. list of the Mlh3 ChIPseq peaks (Excel spreadsheet)**

**Table S3. list of the *pCUP1-IME1* synchronized Spo11 oligonucleotides hotspots (Excel spreadsheet)**

**Table S4:**
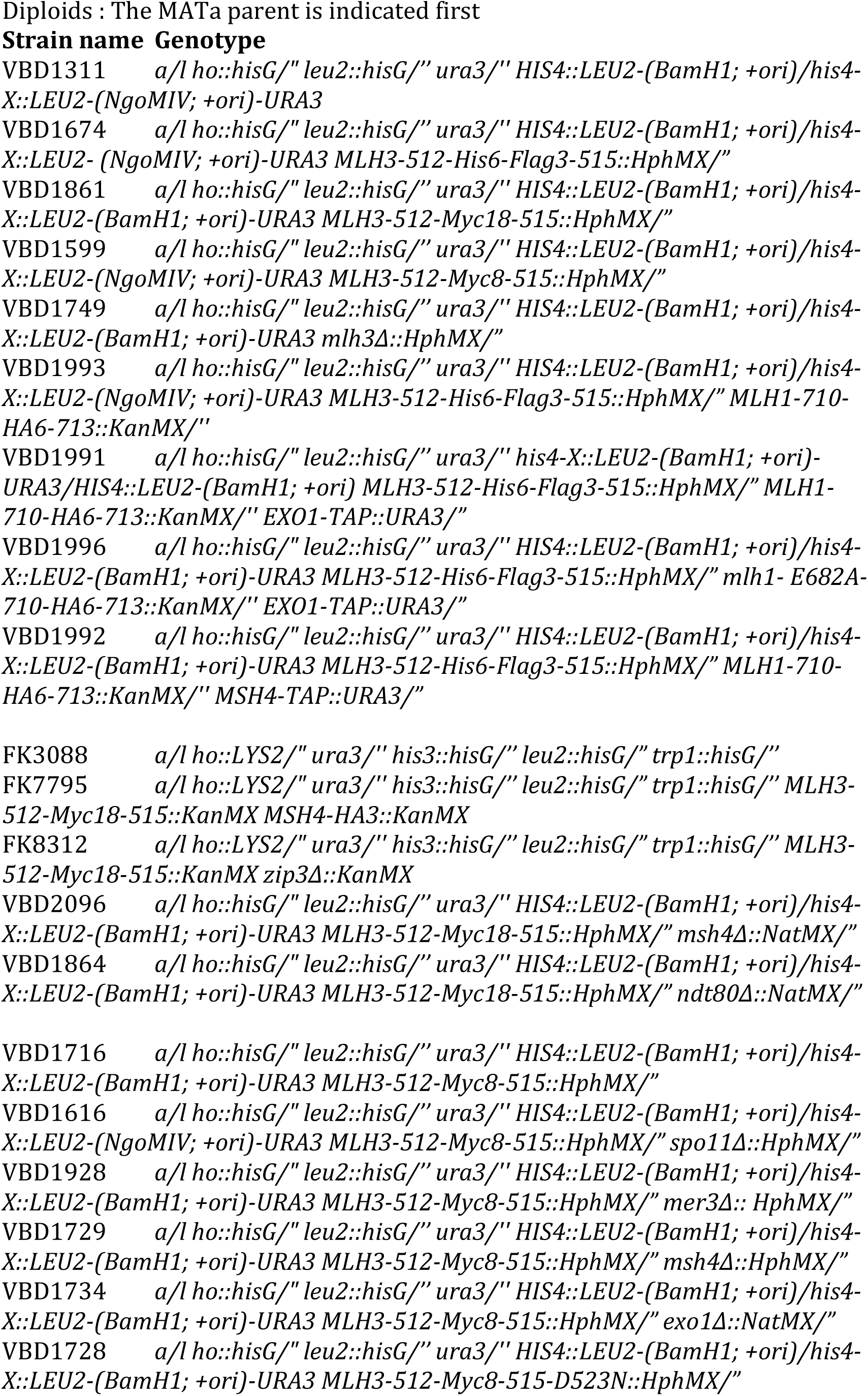
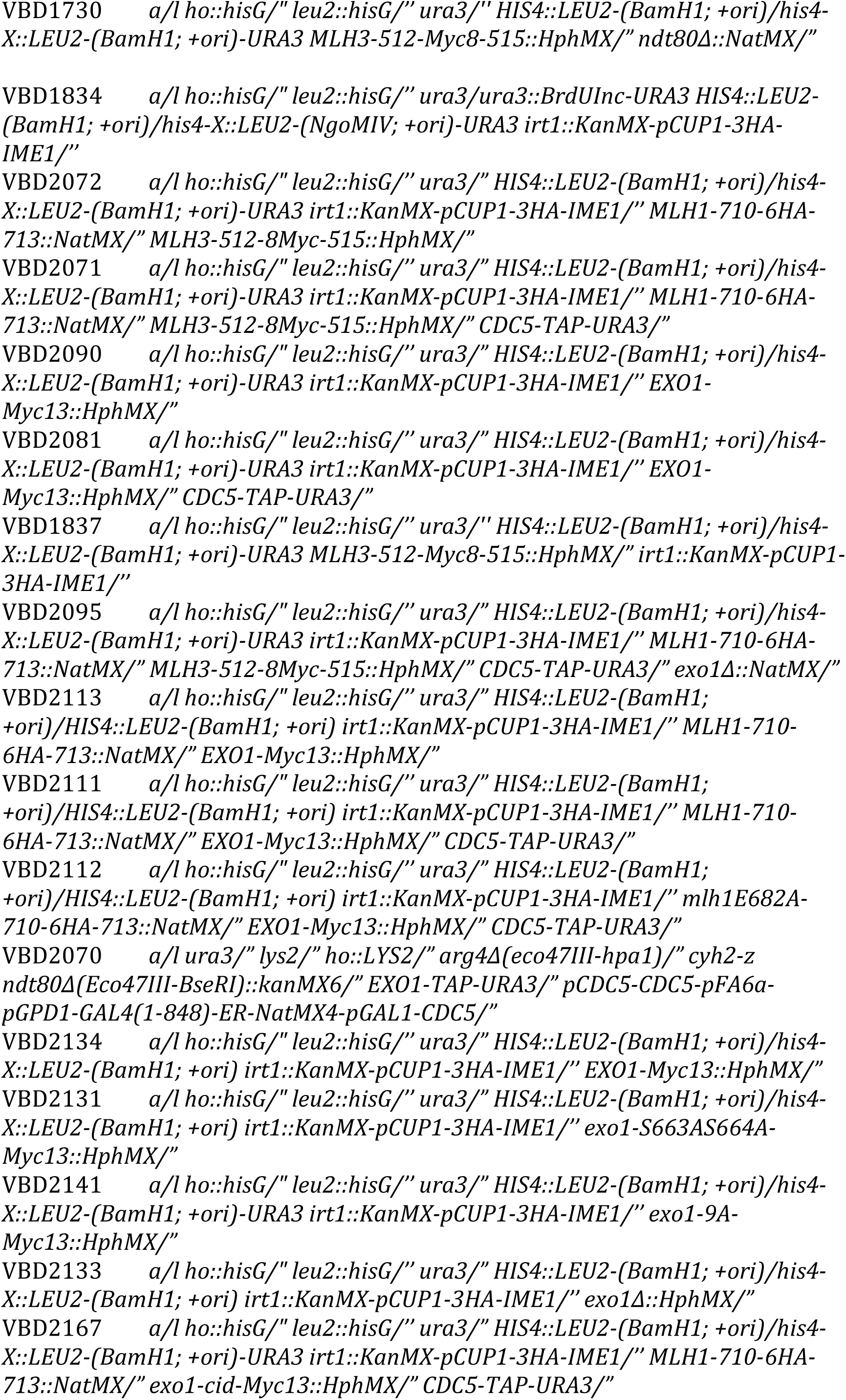
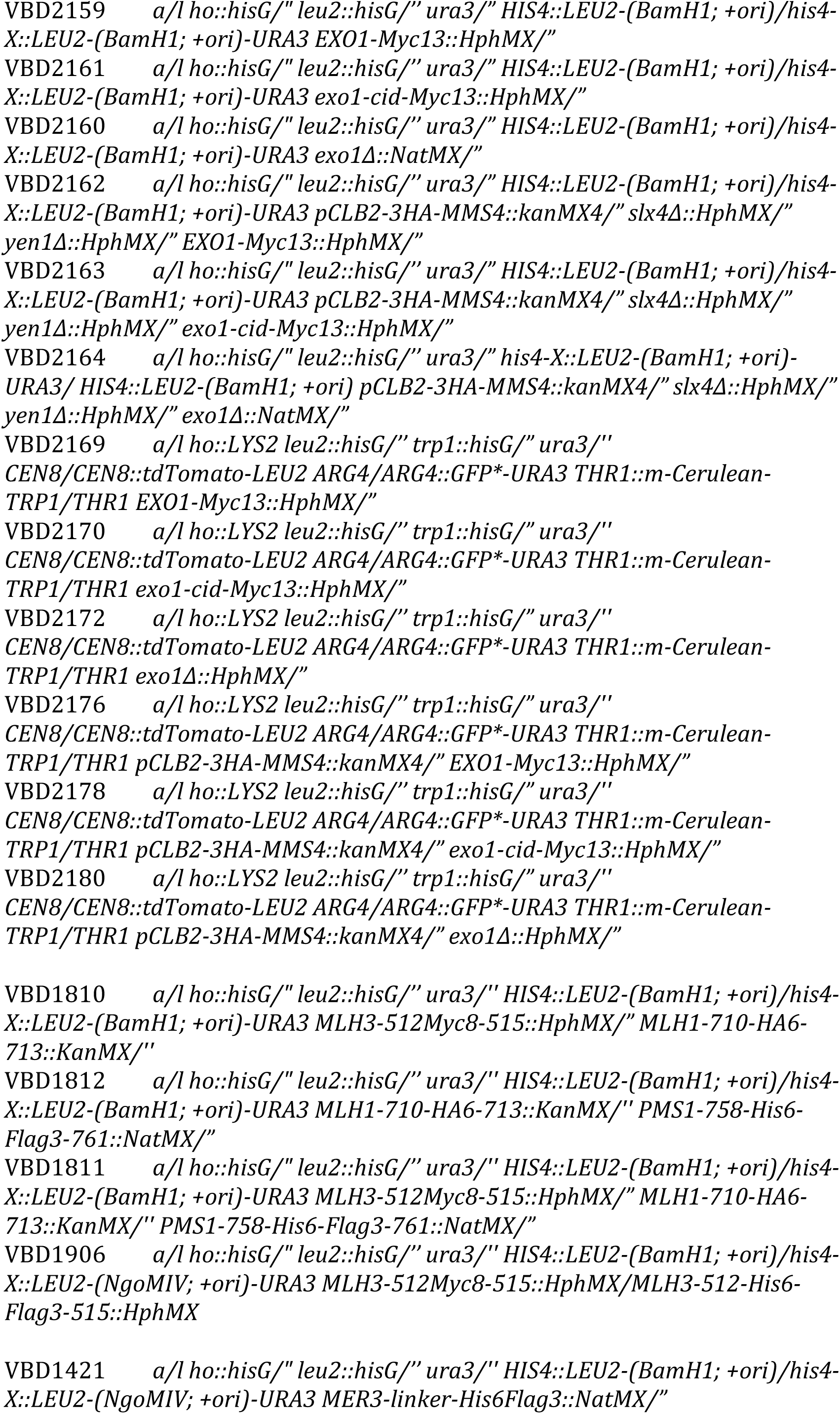
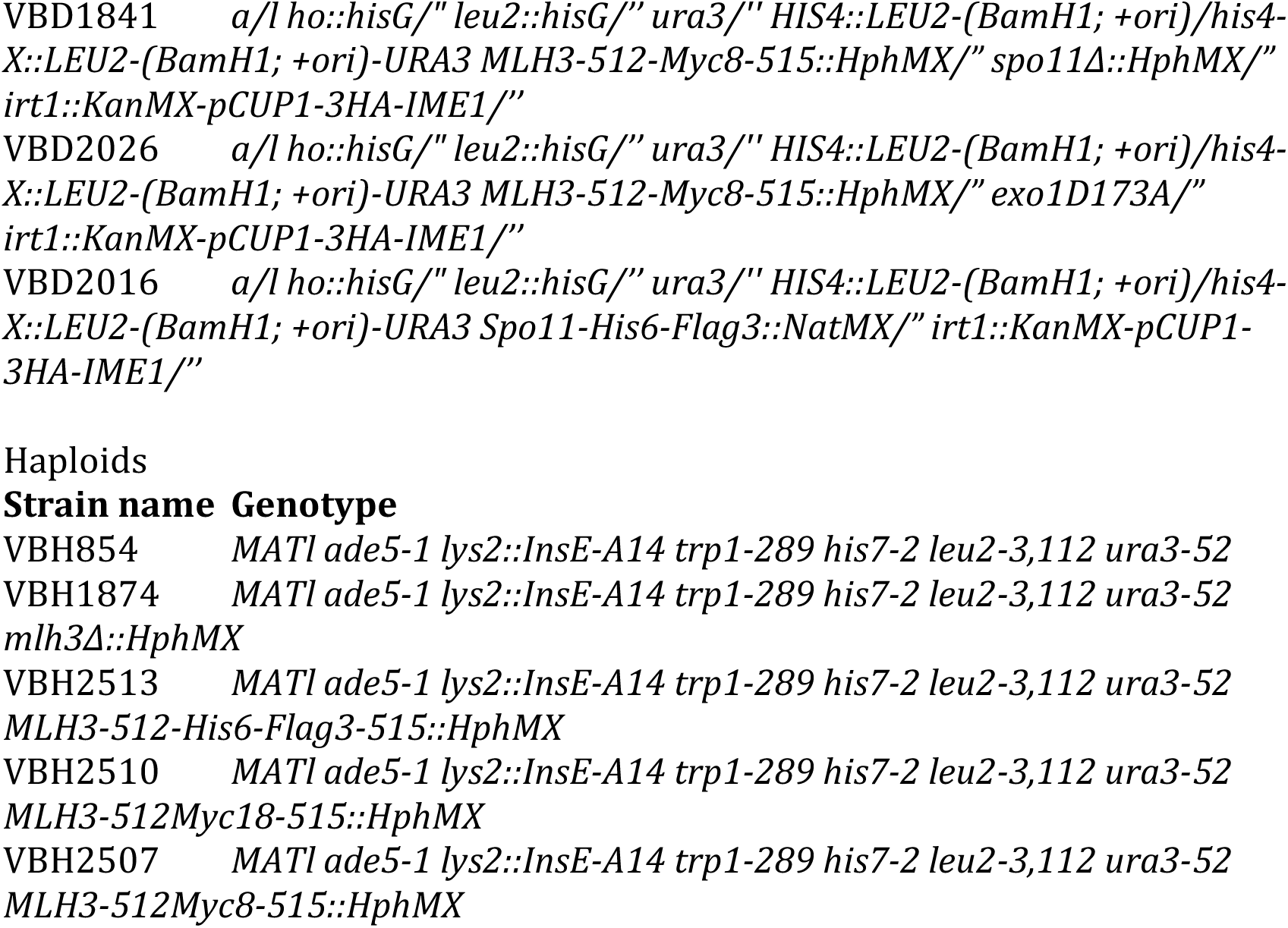
Genotypes of strains used in this study.

**Table S5:**
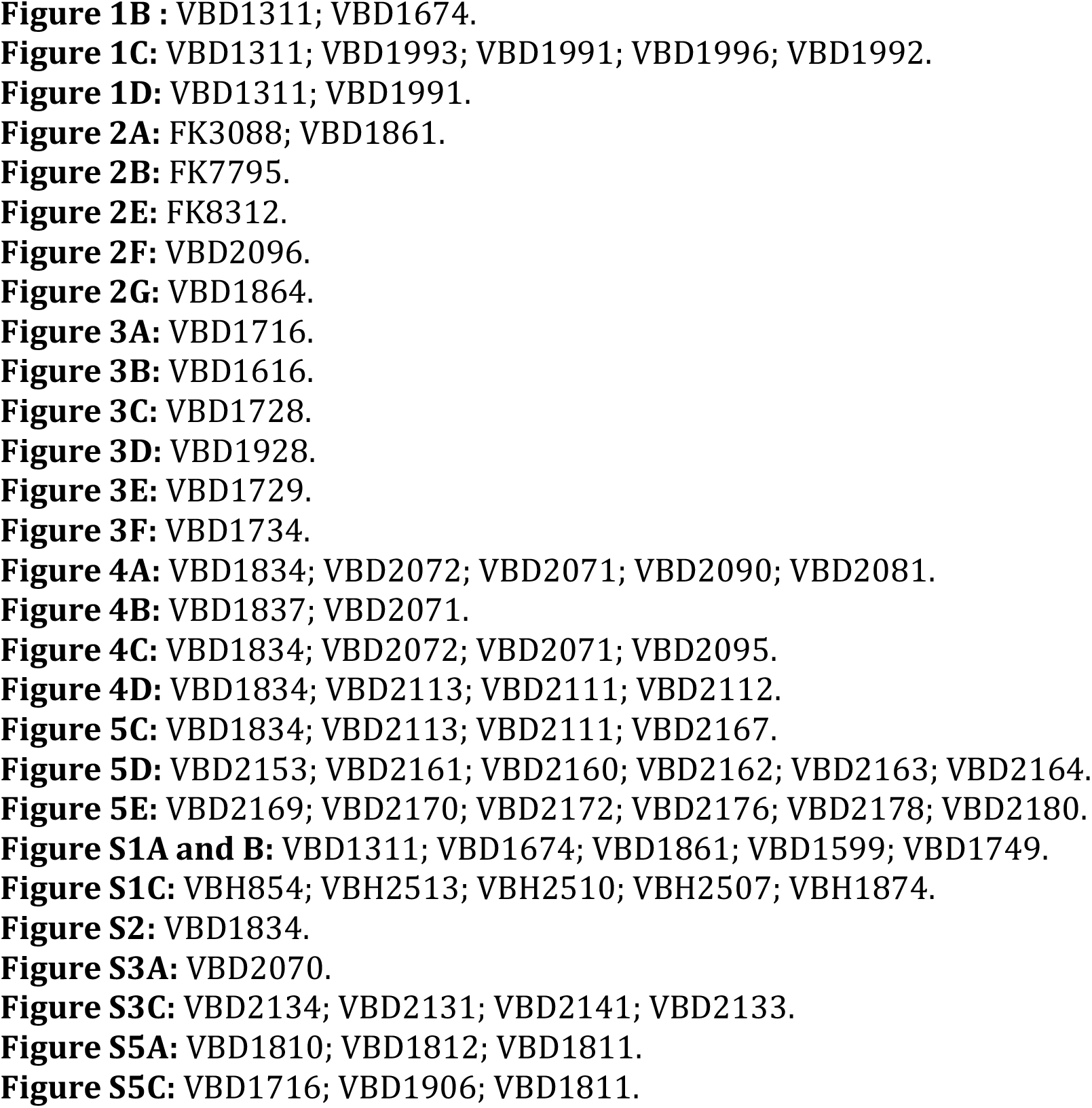
Strains used to the different figures.

**Figure S1.**
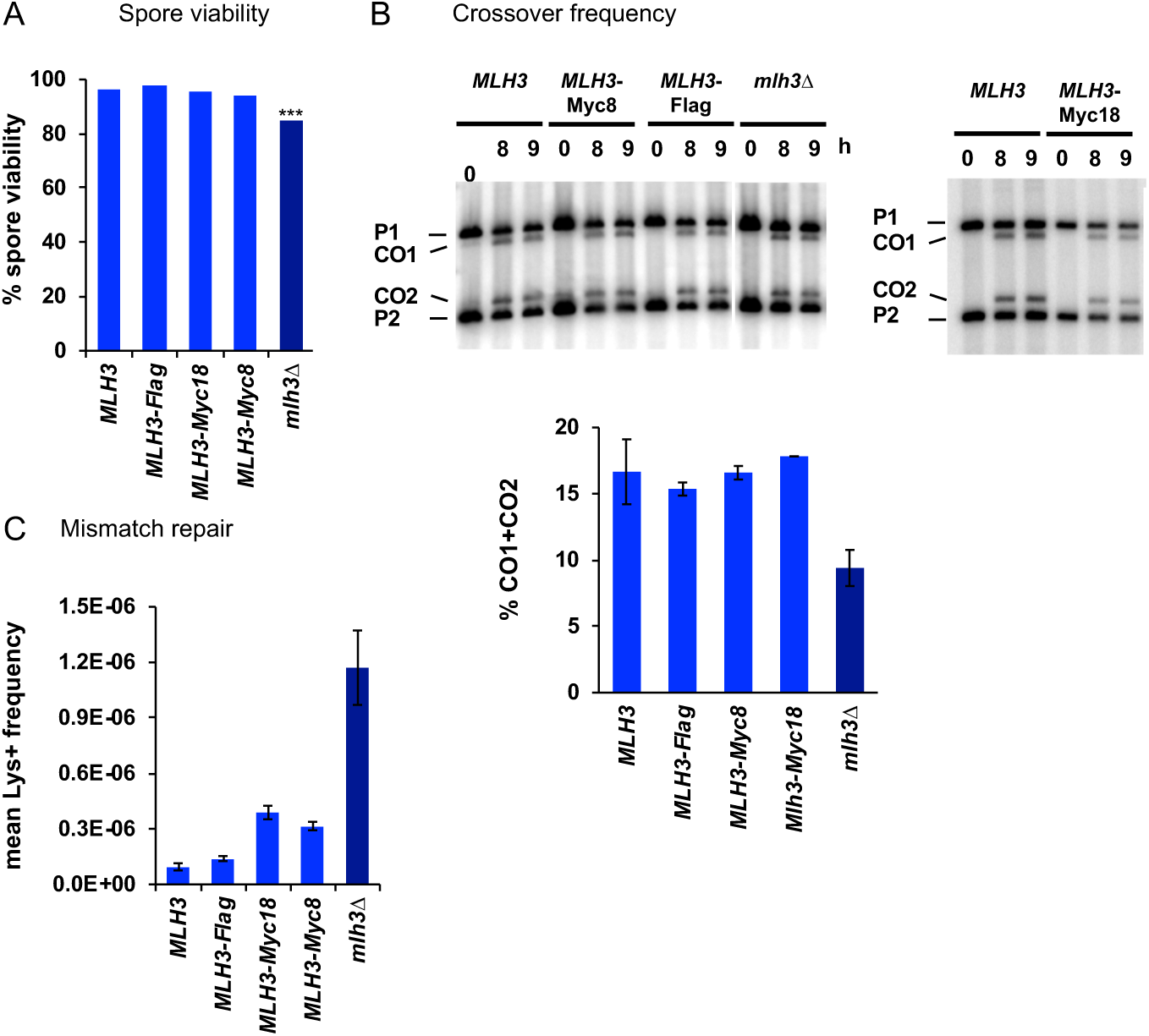
Mlh3 alleles tagged internally in the C terminal domain are functional for meiotic recombination and mismatch repair (A) Spore viability of diploid SK1 strains bearing the indicated *MLH3* genotype at its endogenous locus. *MLH3* (188 tetrads), *MLH3*-Flag (114 tetrads), *MLH3*-Myc18 (104 tetrads), *MLH3*-Myc8 (117 tetrads), *mlh3Δ* (109 tetrads). ***p<4.10^-11^, Fisher’s exact test between MLH3 and *mlh3Δ*. (B) Crossover frequency at the *HIS4LEU2* hotspot monitored by Southern blot at the indicated times in meiosis. Positions of parental bands (P1 and P2) and of the recombinant crossover products (CO1 and CO2) are indicated. Graphs show quantification at 9 h from two (three for MLH3) independent biological replicates ± SD. Same strains as in (A). (C) Mutator assay. Frequency of reversion to Lys+ in haploid vegetative cells containing the indicated *MLH3* genotype at its endogenous locus. Values are the mean ± S.E.M. of 9 independent colonies.

**Figure S2:**
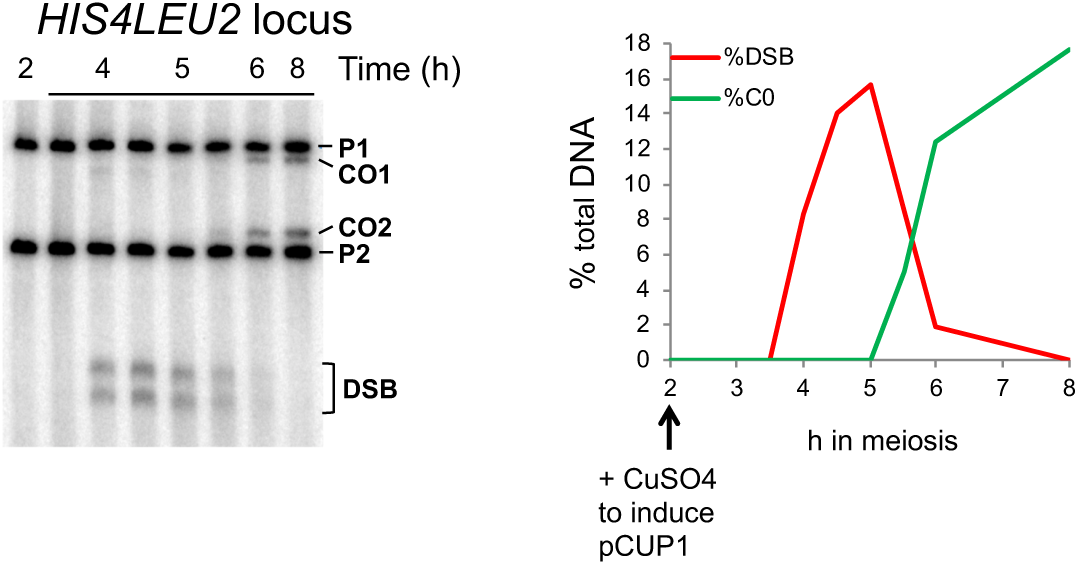
Timing of DSB and crossover formation in p*CUP1-IME1* synchronized meiosis Cu^++^ is added at the 2 h time-point to induce *IME1* expression. DSB and crossover formation is followed at the *HIS4LEU2* hotspot by Southern blot analysis. Positions of parental bands (P1 and P2) and of the recombinant crossover products (CO1 and CO2) are indicated. The graph shows quantification of DSB and crossover frequencies.

**Figure S3:**
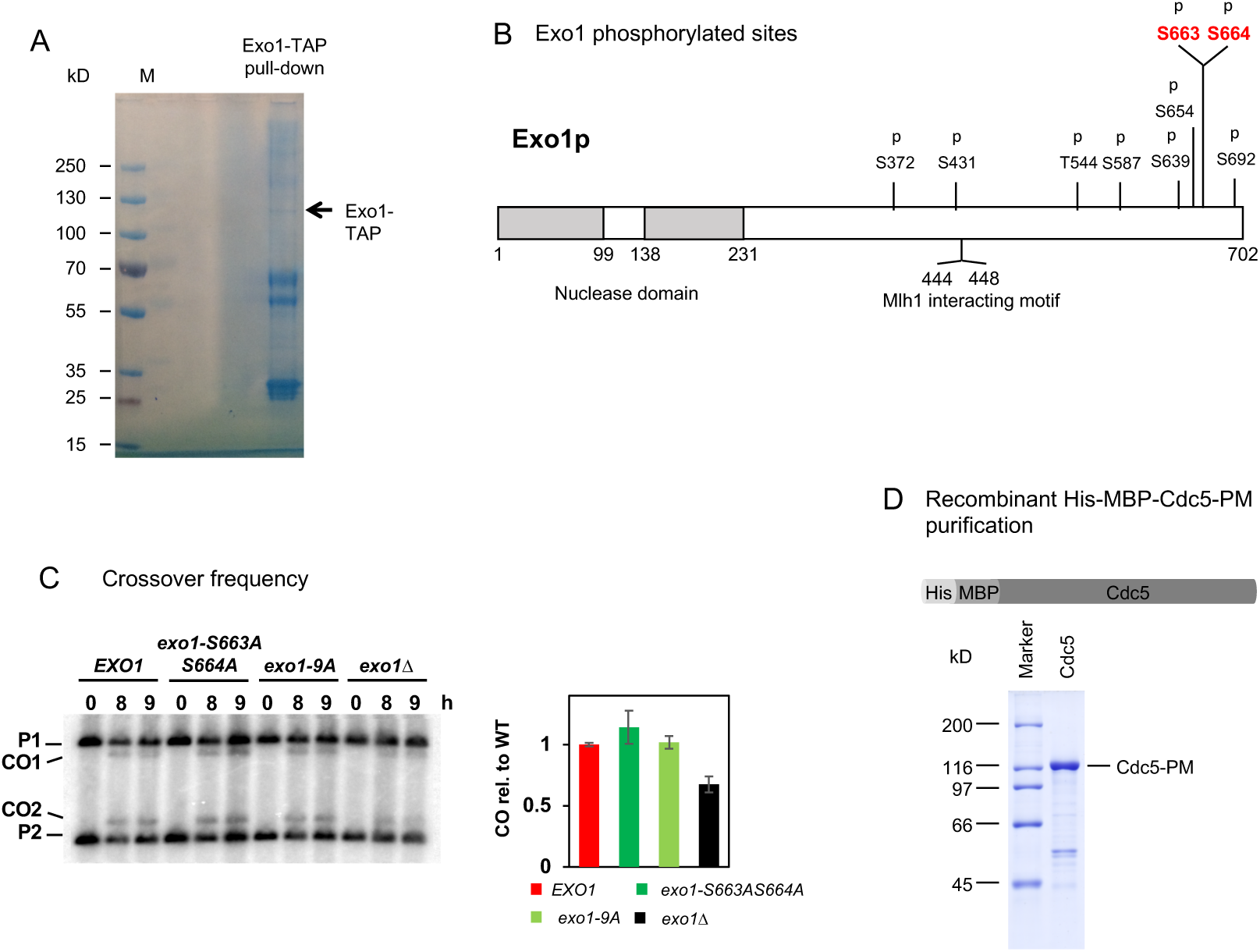
Exo1 phosphorylation sites from cells at the crossover resolution stage of meiosis (A) Colloidal blue stained gel showing the material pulled-down by Exo1-TAP from *ndt80Δ* cells after induction of Cdc5. The band corresponding to Exo1-TAP is indicated with an arrow. (B) Phosphorylated Exo1 residues from the experiment shown in (C). S663 and S664 are within a minimal CDK site and a consensus site for Cdc5 binding (S-S/T-P). Each letter “p” indicates a phosphorylated residue. (C) Crossover frequency at the *HIS4LEU2* hotspot monitored by Southern blot at the indicated times in meiosis. Graph shows quantification at 8 and 9 h ± SD. (D) Purification of recombinant His_MBP-Cdc5-PM (Cdc5 T238D phosphomimetic mutant). Scheme of Cdc5 expression construct. MBP, Maltose binding protein tag; His, 10x histidine tag. The 7.5 % SDS-PAGE shows the final fraction (10 µl of Ni-NTA Eluate) of Cdc5 purification.

**Figure S4:**
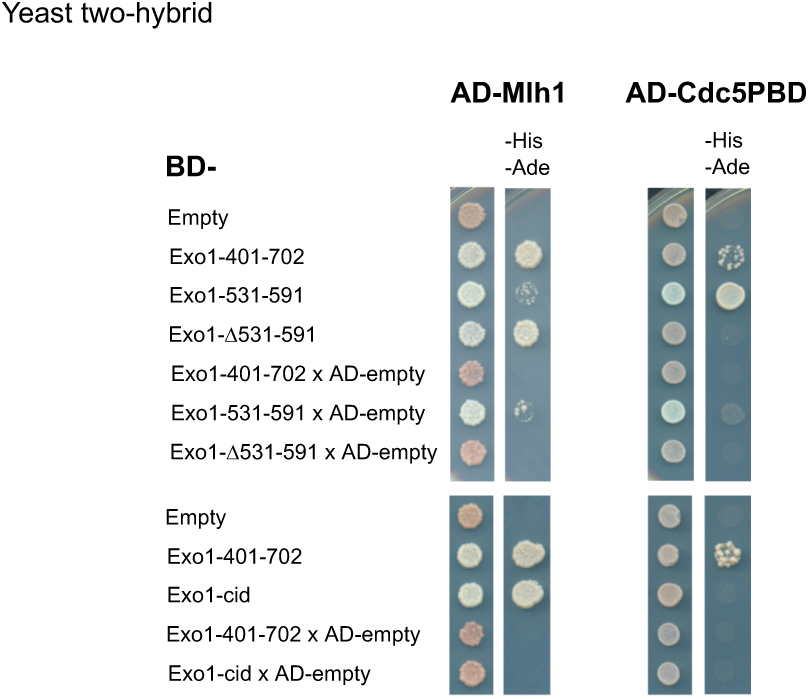
two-hybrid interactions between Exo1 and Cdc5-PBD or Mlh1. The same number of cells of strains expressing the different fusion proteins were plated on minimal media lacking the indicated aminoacids to select for interactions. Growth on -His-Ade medium indicates an interaction. Exo1and Mlh1 interaction is used as a positive control.

**Figure S5:**
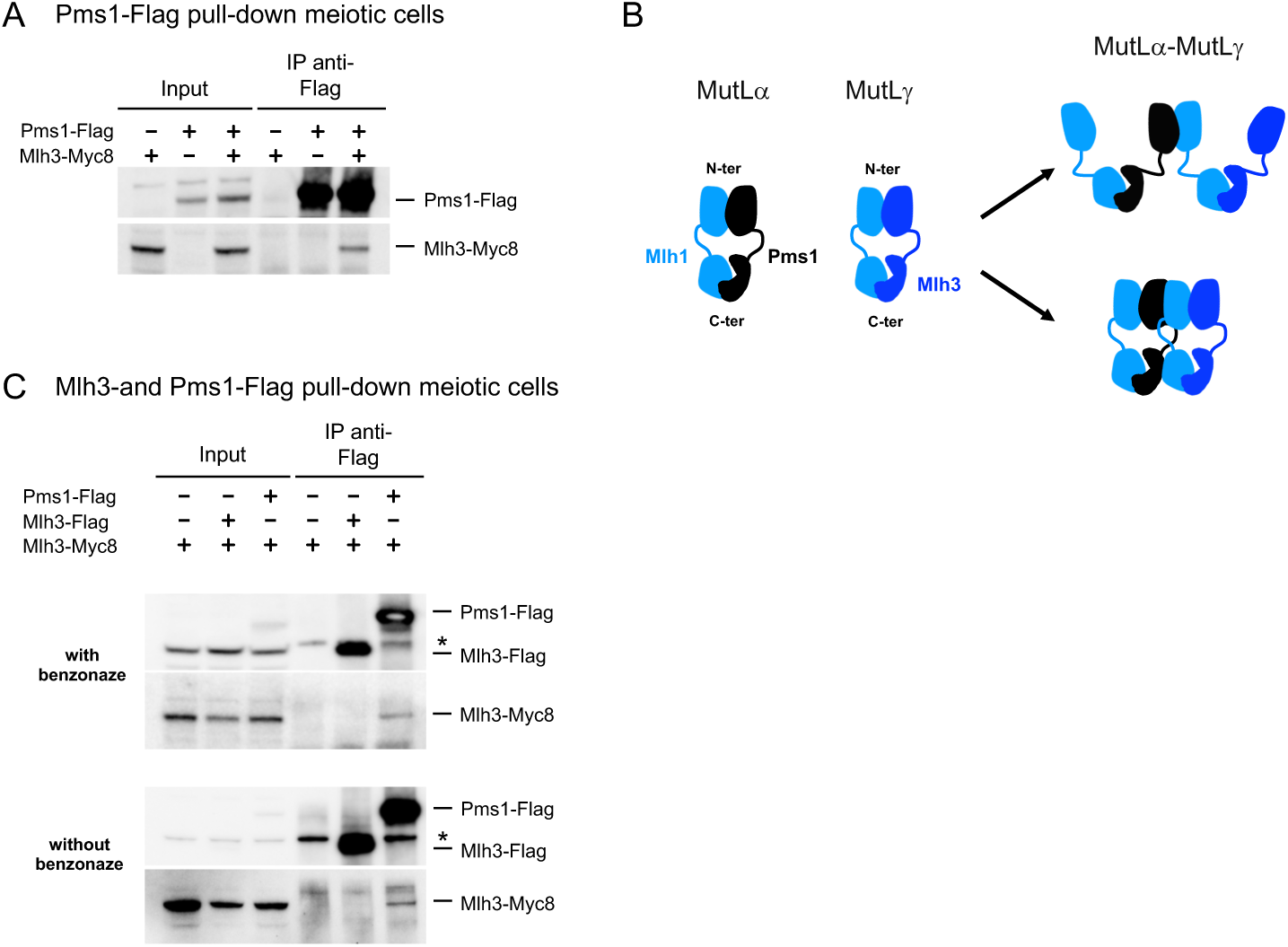
Mlh3 interacts with Pms1 but not with Mlh3 in meiotic cells (A) Coimmunoprecipitation between Pms1-Flag and Mlh3-Myc from cells at 4 h in meiosis after benzonase treatment and analyzed by Western blot. (B) Model for heterotetramer formation between the Nter of Mlh1 and Pms1, or Mlh1 and Mlh3. AFM studies on MutLα^1^ and on MutLγ^2^ showed that these heterodimer can adopt extended conformation with the Cter forming heterodimer and the Nter in the monomer state. We propose that oligomerization of MutLα-MutLγ or of MutLγ-MutLγ could be mediated by interactions of the Nter of two distinct heterodimers. These interactions could be repeated leading to oligomer formation. Such oligomerization could be stabilized when the heterodimers bind DNA. (C) Coimmunoprecipitation between Pms1 and Mlh3 or Mlh3 and itself in meiotic cells at 4 h in meiosis, in the presence (top panel) or absence (bottom panel) of benzonase treatment and analyzed by Western blot.

**Figure S6:**
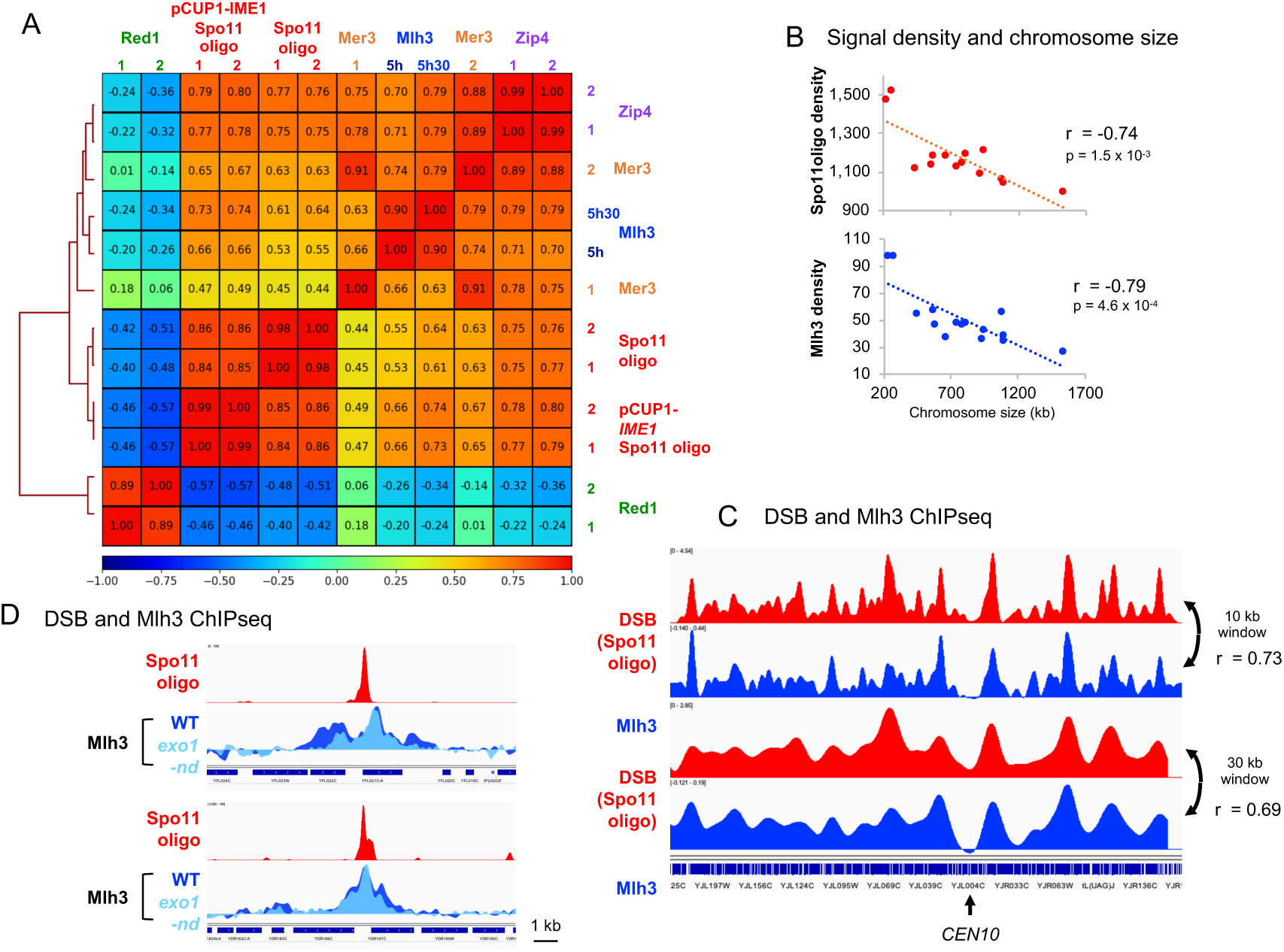
Correlation heatmap between DSBs, Red1, ZMM and Mlh3 binding sites (A) For each replicate, normalized binding data of the indicated protein were used after smoothing with a 2000 bp window. Zip4 data are from^3^, Spo11 oligo data are from^4^ and Red1 data are from^5^. The comparison was made on the regions encompassing the Red1 peaks (933 peaks)^5^, the top 1000 Spo11 oligo hotspots^4^ and the top 1000 Mlh3 peaks determined at 5.5 h (see Table S2). The Spearman correlation coefficient is indicated for each pair-wise comparison. (B) Like DSBs, Mlh3 density is stronger on small chromosomes. The total signal of Spo11oligonucleotides from a p*CUP1-IME1* time-course or the Mlh3 ChIPseq at 5.5 h per chromosome is plotted as a function of chromosome size. Chromosome III was excluded from the analysis, due to the presence of the *HIS4LEU2* hotspot in the strains used. (C) Chromosome-wide correlation between DSB (p*CUP1-IME1* Spo11 oligos) and Mlh3, at two different smoothing scales. The correlation coefficient (Spearman) along the whole genome is indicated for each comparison. (D) Mlh3 ChIPseq signal in WT or *exo1-nd* mutant at two Spo11 oligo hostpots. Spo11 oligo profiles are from a p*CUP1-IME1* synchronized time-course. Signal is smoothed with a 200 bp window.

**Figure S7:**
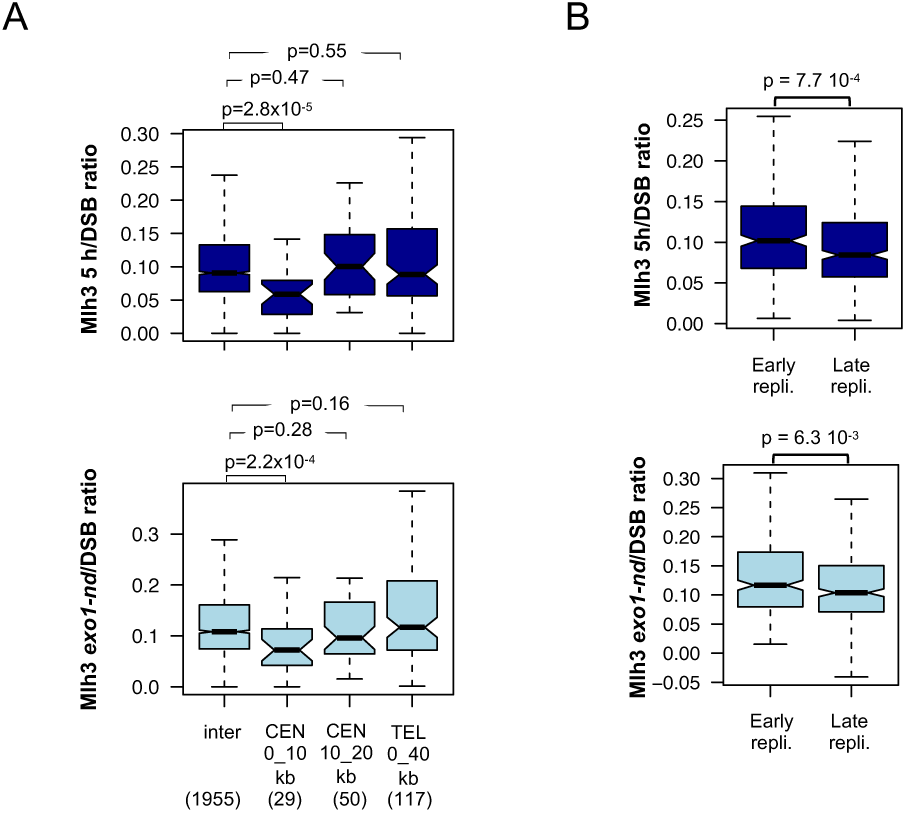
Mlh3 signals per DSB in proximity to a centromere or a telomere and in early versus late replicated regions The normalized ChIPseq signal of each protein divided by the corresponding p*CUP1-IME1* Spo11 oligo was computed on the width plus 1 kb on each side of the strongest 2000 hotspots^4^ at the indicated chromosome regions. (A) inter, CEN 0-10kb, CEN 10-20kb and TEL 0-40 kb regions as in Figure 6C. (B) Early and late replicating regions as in Figure 6E. Data are represented as boxplots, and the statistical differences (Mann-Whitney-Wilcoxon test) between different regions are indicated.

## SUPPLEMENTAL MATERIALS AND METHODS

### Mutation analysis

Mutation rates in the presence of tagged *MLH3* alleles were estimated by measuring the spontaneous reversion rate at the *lys2::InsE-A14* locus in strains derived from the E134 strain^6,7^. Three single colonies from three independent transformants of the same genotype or nine colonies of the parent E134 strain were grown to stationary phase in liquid YPD medium and plated onto YPD or selective medium lacking lysine for revertant count.

### Exo1-TAP pull down and phosphopeptides analysis

*ndt80Δ* cells containing an inducible *CDC5* gene were used^8^. After SPS pre-growth and 7 h in sporulation medium, 1 µM Estradiol was added to induce p*GAL1*-*CDC5*. 2.10^10^ cells were harvested after 8 h 15 min in sporulation medium and processed as described above for the Flag-affinity pull-down in the presence of 125 U/mL benzonase. 1500 μL of PanMouse IgG magnetic beads (Thermo Scientific) were washed 1:1 with lysis buffer, preincubated in 100 μg/ml BSA in lysis buffer for 2 h at 4°C and then washed twice with 1:1 lysis buffer. The lysate was cleared by centrifugation at 8000 g for 10 min at 4°C and incubated overnight at 4°C with the washed PanMouse IgG magnetic beads. The magnetic beads were washed four times with 8 mL of wash buffer (20 mM HEPES/KOH pH7.5; 150 mM NaCl; 0.5% Triton X-100; 5% Glycerol; 1 mM MgCl2; 2 mM EDTA; 1 mM PMSF; 1X Complete Mini EDTA-Free (Roche)). Beads were then resuspended in 40 μl of Laemmli buffer and boiled for 10 min at 75°C. Proteins were separated by SDS-PAGE, stained with colloidal blue, and the band predicted to contain Exo1-TAP was excised and processed. Excised gel slice was washed and proteins were reduced with 10 mM DTT prior to alkylation with 55 mM iodoacetamide. After washing and shrinking of the gel pieces with 100% acetonitrile, in-gel digestion was performed using trypsin/LysC (0.1µg) overnight in 25 mM ammonium bicarbonate at 30°C. Peptide are then extracted using 60/35/5 MeCN/H2O/HCOOH, vacuum concentrated to dryness and reconstituted in loading buffer A (2/98 MeCN/H2O + 0.05% TFA) prior to liquid chromatography-tandem mass spectrometry (LC-MS/MS) analysis. The extracted peptides were chromatographically separated using an RSLCnano system (Ultimate 3000, Thermo Scientific) coupled to a

Q Exactive HF-X mass spectrometer (Thermo Scientific). For identification the data were searched against the UniProtKB/Swiss-Prot *S. cerevisiae* database using Sequest with Proteome Discoverer (version 2.2) and SwissProt fasta database containing *S. cerevisiae* sequences using MascotTM (version 2.5.1). The resulting files were further processed using myProMS^9^ v3.6. The maximum false discovery rate (FDR) calculation was set to 1% at the peptide level for the whole study (Percolator or QVALITY algorithm). We validate phosphorylated peptides by combining the phosphoRS informations and by manually inspecting the peak assignment. Data are available via ProteomeXchange with the identifier PXDO14185.

